# Integration and segregation in Autism Spectrum Disorders modulated by age, disease, and interaction: A graph theoretic study of intrinsic functional connectivity

**DOI:** 10.1101/278846

**Authors:** Vatika Harlalka, Shruti Naik, Raju S. Bapi, P.K. Vinod, Dipanjan Roy

## Abstract

Autism spectrum disorder (ASD) is a neurodevelopmental disorder affecting 1 in 50 children between the ages of 6 and 17 years. Brain connectivity and graph theoretic methods have been particularly very useful in shedding light on the differences between high functioning autistic children compared to typically developing (TD) ones. However, very recent developments in network measures raise a cautionary note by highlighting gross under- and over-connectivity in ASD may be an oversimplified hypothesis. Thus the primary aim of our study is to investigate these notions in functional connectomics of ASD versus TD by subjecting the data to reproducibility experiments using two independent datasets.

Further, we tested the hypothesis of alteration in network segregation and integration in the ASD subjects. We have analyzed the resting-state functional magnetic resonance imaging (rs-fMRI) and diffusion tensor imaging (DTI) data from the University of California Los Angeles (UCLA) multimodal connectivity database (n=42 ASD, n=37 TD) and rs-fMRI data from the Autism Brain Imaging Data Exchange (ABIDE) (n=187 ASD, n=176 TD) dataset. We assessed the differences in connection strength between TD and ASD subjects. We also performed graph theoretical analysis to analyze the effect of disease on various network measures. Further, using the larger ABIDE dataset, we performed two-factor ANOVA test, to study the effect of age, disease and their interaction by classifying the TD and ASD participants into two cohorts: children (9-12 years, n=73 TD and n=87 ASD) and adolescents (13-16 years, n=103 TD and n=100 ASD). In ASD, we show the existence of atypical connectivity within and between functional networks as compared to TD. We also found in ASD both hypo-and hyper-connectivity within functional networks such as the default mode network (DMN). Further, graph theoretic analysis showed that there is significant effect of age and disease on modularity, clustering coefficient, and local efficiency. We also identified specific areas within the DMN, sensorimotor, visual and attention networks that are affected by age, disease and their interaction. Overall, our findings suggest that maturation, disease and their interaction are critical for unraveling the biological basis and developmental trajectory in ASD and other neuropsychiatric disorders.

## Introduction

Autism spectrum disorder (ASD) is a clinical umbrella term for neurodevelopmental disorders (Kim et al., 2011; Zablotsky et al.; 2014) that are characterized by atypical social behavior, including deficits in receptive and expressive language, theory of mind (TOM), and cognitive flexibility deficit. It encompasses autism, Asperger syndrome, pervasive developmental disorder not otherwise specified (PDD-NOS), and childhood dis-integrative disorder. Accumulating evidence indicates that ASD is associated with alterations of neural circuitry compared to typically developing (TD) individuals (Di Martino et al., 2009; McAlonan et al., 2005; Schmitz et al., 2006; Stanfield et al., 2008; Vissers et al., 2012; Rudy et al., 2013; Di Martino et al., 2013; Supekar et al., 2013; Keown et al., 2013). However, the recent literature is divided as to whether the Autistic brain is dominated by under- or over-connectivity among the structural and functional brain regions. A method of choice to study the brain connectivity and its deviation in disease from the healthy controls has been the resting state functional magnetic resonance imaging (rs-fMRI), which detects correlated fluctuations in the blood level oxygen dependent (BOLD) signals from the brain regions giving rise to what is known as the resting state functional connectivity (rs-FC). While the majority of the early findings in rs-FC suggested under-connectivity in midline core in the proximity of default mode brain regions in ASD compared to TD (Kennedy and Courchesne, 2008; Vissers et al., 2012; Yerys, Benjamin E. et al., 2015), a growing number of studies with advanced methodological details (e.g. Nair et al., 2013; Salmi et al., 2013; Hahamy et al., 2015; Uddin et al., 2015) suggest a rather complex picture that also includes over-connectivity in children with Autism (Rudie et al., 2013; Uddin et al., 2015; Keown et al., 2017), possibly related to impaired network differentiation (Rudie et al., 2012; Fishman et al., 2014; Fishman et al., 2015). Recently, the focus has shifted to looking at the hypothesis of topographic distribution of connectivity patterns (Hahamy et al., 2015; Uddin et al., 2015).

Further, studies also suggest a reduced connectivity pattern within major networks (functional segregation) and increased connectivity between different networks (functional integration) in ASD (Rudie et al., 2011; 2012). Functional brain networks become simultaneously more integrated and segregated during typical development (e.g., Fair et al., 2009) and white matter integrity increases during development (e.g., Lebel et al., 2012), suggesting that brain networks in ASD may reflect ‘immature’ or aberrant developmental processes. Despite this array of regional and systems-level findings in ASD, it is still unclear how these alterations might be reflected at a network-level where the brain is modeled as a network of hundreds of interacting and competing regions of several integrated and segregated systems. Graph theory, which describes complex systems as a set of “nodes” (i.e., brain regions) and “edges” (i.e., connections between nodes), has characterized the brain as a complex network with a hierarchical modular organization consisting of several major functional communities (i.e., visual, sensorimotor, default mode, and attentional systems; see Wang et al., 2010, for review). Structural and functional brain networks exhibit robust levels of local and global efficiency (i.e., small-world properties; Watts and Strogatz, 1998) that can be quantitatively characterized using graph theoretic methods (Bullmore and Sporns, 2009; Rubinov and Sporns, 2010). Given the complexities and inconsistencies of the ASD literature, graph theoretic data-driven techniques provide exploratory approaches that are suitable for uncovering connectivity patterns even in the absence of strong directional hypotheses. Using graph theoretical analysis on rs-FC, studies have reported a decrease in local efficiency, clustering coefficient and characteristic path length, and an increase in global efficiency, in the ASD subjects, which is interpreted as increased randomness of functional network organization (Rudie et al., 2012; Itahashi et al., 2014). Interestingly, recent studies observed atypically increased intrinsic functional connectivity (iFC) inside the “rich club” (i.e., densely interconnected hubs) in ASD, both in a small in-house sample and in a larger low-motion multi-site data set selected from the ABIDE, and age associated change in the “rich club” organization using UCLA Multimodal Connectivity Database (Rudie et al., 2012).

In this study, we analyzed the functional magnetic resonance imaging (fMRI) and diffusion tensor imaging (DTI) data from the University of California Los Angeles (UCLA) multimodal connectivity database and functional MRI data from the Autism Brain Imaging Data Exchange (ABIDE) dataset to reproduce some of the results related to the functional connectomics of ASD versus TD and to test the altered functional integration and segregation hypothesis in ASD. To do this, we first studied the distribution of atypical connectivity over the whole brain and within communities. We assessed the differences in connection strength and computed various local and global graph theoretic metrics such as modularity, characteristic path length, small-worldness, local and global efficiency. Using these measures, we assessed the differences of functional integration and segregation between TD and ASD. We analyzed the change in the network measures in younger children as compared to adolescents by dividing our dataset into two cohorts based on age and performed 2 × 2 ANOVA analysis to study the effect of age, disease and their interaction on network measures.

## Materials and Methods

### Data

The data included in the current study were selected from ABIDE Preprocessed Initiative (Di Martino et al., 2014; Craddock et al., 2013) and UCLA multimodal connectivity database acquired as a part of the human connectome project (HCP) (Brown et al., 2012). A subsample from the ABIDE fMRI dataset was considered that comprises two cohorts: children (9-12 years, 73 TD and 87 ASD) and adolescent (13-16 years, 103 TD and 100 ASD). ABIDE dataset used for this study was collected from twenty different centers. Institutional review board approval was provided by each data contributor in the ABIDE database. Detailed recruitment and assessment protocols and inclusion criteria are available on the ABIDE website. The ABIDE data set was made public in August 2012 and can be accessed from: http://fcon_1000.projects.nitrc.org/indi/abide/. The preprocessed data is available at: http://preprocessed-connectomes-project.org/abide/. The UCLA multimodal connectivity database contains fMRI data for 37 TD (mean age: 13+/-2, age range: 9.5–17.8) and 42 ASD (13+/-2.4, age-range 9.3–17.9) subjects, which was used for our functional connectivity analysis. On the other hand, for the whole brain structural connectivity, UCLA DTI dataset consisting of 43 TD (13.1 +/-2.4, 9.0–18.0) and 51 ASD (13.0 +/-2.8, 8.4–18.2) subjects was used. Informed consent and assent to participate was obtained prior to assessment according to protocols approved by the UCLA Institutional Review Board (IRB). The consent obtained for both UCLA and ABIDE datasets were informed and written. The parents and the legal guardians of children and adolescents (non-adult participants) provided written informed consent. Participant details are provided in the **Tables 1 and 2** (also see Supplementary Tables **S1** and **S2** for more information).

**Table 1:**
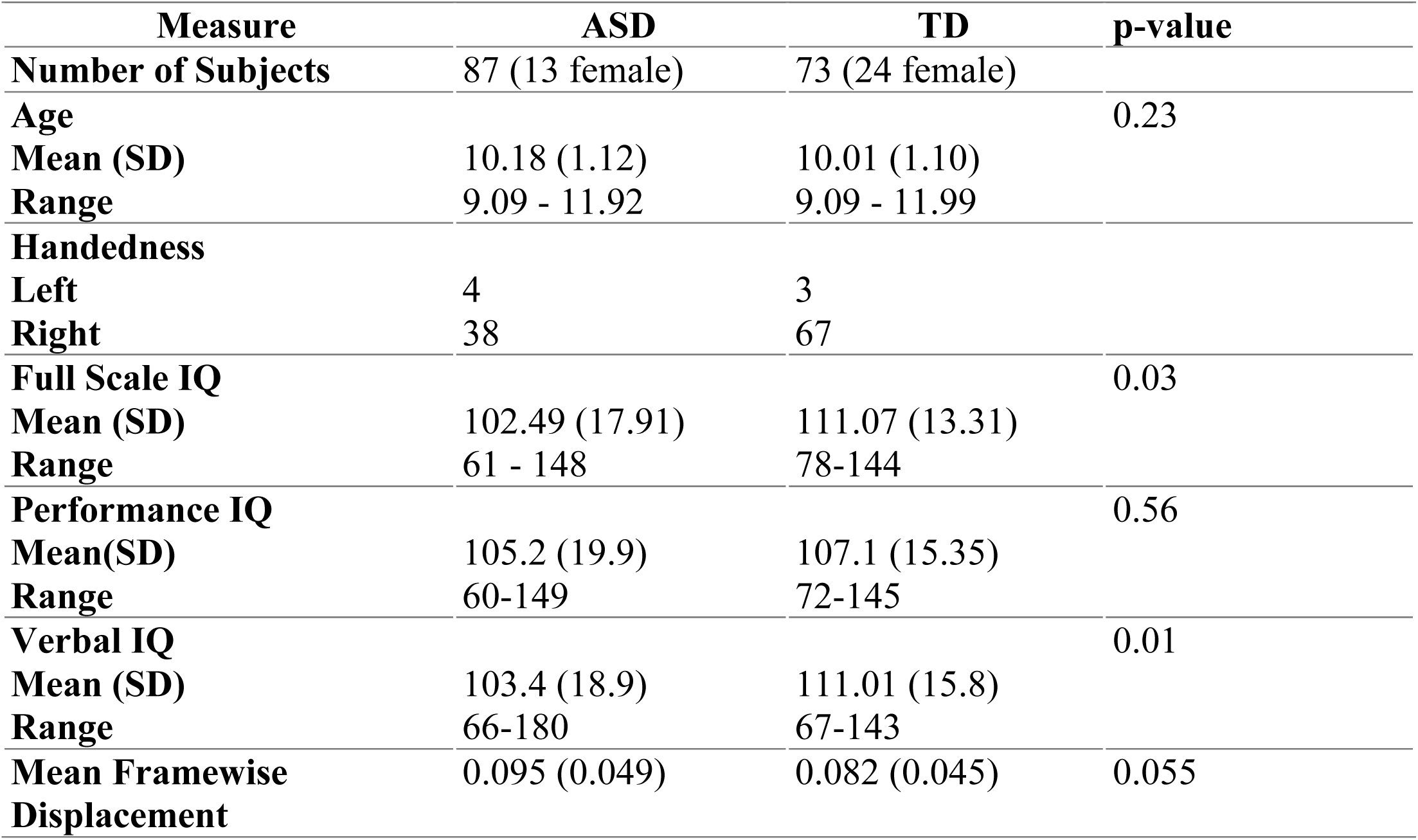
Participant information of a subset of ABIDE dataset used for the analysis in the age group 9-12 years.

**Table 2:** Participant information of a subset of ABIDE dataset used for the analysis in the age group 13-16 years* (*Handedness data were missing for 77 ASD and 25 TD).

| Measure | ASD | TD | p-value |
| --- | --- | --- | --- |
| **Number of Subjects** | 100 (10 female) | 103 (30 female) |  |
| **Age** |  |  | 0.32 |
| **Mean (SD)** | 14.35 (0.80) | 14.50 (0.79) |  |
| **Range** | 13.04 - 15.9 | 13.1 - 15.95 |  |
| **Handedness** |  |  |  |
| **Left** | 11 | 10 |  |
| **Right** | 57 | 71 |  |
| **Full Scale IQ** |  |  | 0.27 |
| **Mean (SD)** | 105.58 (16.3) | 108.01 (13.31) |  |
| **Range** | 75-139 | 73-138 |  |
| **Performance IQ** |  |  | 0.87 |
| **Mean(SD)** | 105.0 (16.48) | 105.32 (13.51) |  |
| **Range** | 59-140 | 67-139 |  |
| **Verbal IQ** |  |  | 0.02 |
| **Mean (SD)** | 102.42 (18.3) | 109.6 (13.6) |  |
| **Range** | 62-141 | 73-140 |  |
| **Mean Framewise Displacement** | 0.081 (0.04) | 0.071 (0.035) | 0.098 |

### Data Acquisition and Preprocessing

For the sake of completeness, in the following sections, we summarize the details of acquisition protocols followed to obtain UCLA and ABIDE data sets.

### UCLA HCP Dataset

All the resting-state fMRI and DTI scans were acquired on a Siemens 3T Trio at UCLA. Subjects were asked to keep their eyes open and attend to a fixation cross displayed for 6 min (T2*-weighted functional images: TR = 3000 ms, TE = 28 ms, matrix size 64 × 64, 19.2 cm FoV, and 34 4-mm thick slices (no gap), interleaved acquisition, with an in-plane voxel dimension of 3.0×3.0 mm). The DTI sequence included 32 directions (*b* = 1000 s/mm^2^), three scans with no diffusion sensitization, at *b* = 0, and additional six scans at *b* = 50 s/mm^2^. Other parameters were TR = 9500 ms, TE = 87 ms, GRAPPA on, FOV = 256 mm, with 75 axial slices of 2×2×2 mm^3^). Other details about the acquisition parameters and protocol can be found in Rudie et al. (2012).

Functional and structural connectivity matrices were calculated using the 264 ROI functional parcellation scheme which was adopted from a meta-analysis carried out recently by Power et al. (2011). This has provided a comprehensive subnetwork identification of functionally specialized regions of interest based on large number of fMRI samples. For each subject, aggregated BOLD time-series of the 264 brain regions were used to compute pairwise Pearson correlation coefficients which were then z-transformed to obtain a 264 × 264 whole brain functional connectivity matrix. DTI tractography was carried out by propagating fibers from each voxel without a Fractional Anisotropy (FA) cutoff. A fiber was defined as connecting two regions if one fiber endpoint terminated within one region and the other endpoint terminated within the other region. This process was repeated using all the 264 regions as seeds to derive a 264 × 264 whole brain structural connectivity matrix for each subject (Rudie et al., 2013).

### ABIDE Dataset

Data acquisition was performed at a magnetic field strength of 3.0 Tesla (Magnetom VERIO, Siemens, Erlangen, Germany). During the 6-minute resting state functional experiment participants were instructed to minimize head movement and stay awake with their eyes closed. Data was acquired using T2*-weighted echo planar imaging (TE = 30 ms, TR = 3000 ms, flip angle = 90°, spatial resolution 3×3×4 mm^3^, 36 slices, 120 volumes). For anatomical reference a high-resolution isotropic Magnetization Prepared Rapid Gradient Echo (MPRAGE) sequence was acquired (TE = 7.6 ms, TR = 14 ms, flip angle = 20°, spatial resolution 0.8×0.8×0.8 mm^3^, 160 slices). All high-resolution T1 MPRAGE images were quality controlled by a board-certified radiologist to detect potential artifacts such as ringing due to motion, spiking or signal loss. The details of image acquisition described above have been obtained from the ABIDE website.

ABIDE resting-state fMRI data was preprocessed using the configurable pipeline for the analysis of connectomes (CPAC) with bandpass filtering. CPAC preprocessing includes slice time correction, motion correction, skull strip, global mean intensity normalization and nuisance signal regression. The details can be found here: http://preprocessed-connectomes-project.org/abide/cpac.html. Any scans exhibiting artefacts, signal dropout, suboptimal registration or excessive motion were excluded. Further, we only used the quality-assured scans from the ABIDE Preprocessed Initiative open database. Functional parcellation was accomplished using the two-stage spatially-constrained functional procedure (Craddock et al., 2012, Craddock et al., 2013). Consensus regions between Craddock-200 and Power-264 parcellations were identified to align the region labels in the Craddock-200 parcellation (see **Supplementary Table S3 and S4**). Functional connectivity is calculated using the correlations between the BOLD time-series from the 200 regions of the brain defined in the Craddock atlas (Craddock et al., 2012). To do this, aggregated BOLD time-series of each area was used to calculate the pairwise Pearson correlation coefficients which were then z-transformed to obtain a 200 × 200 functional connectivity matrix for each subject in TD and ASD. Subsequently, the mean connectivity matrix was calculated for TD and ASD children and adolescent subgroups.

### Graph Theoretic Analysis Methods

The workflow of the graph theoretic analysis on the datasets is depicted in **Figure 1**. We used data obtained from UCLA multimodal database to study the disease effect on structural and functional connectivity. To further analyze the functional connectivity, we considered the larger ABIDE dataset where we could age-stratify the data to look at the age, disease and their interaction effects on the various network measures. The details of the methods are given the following sections.

**Fig. 1:**
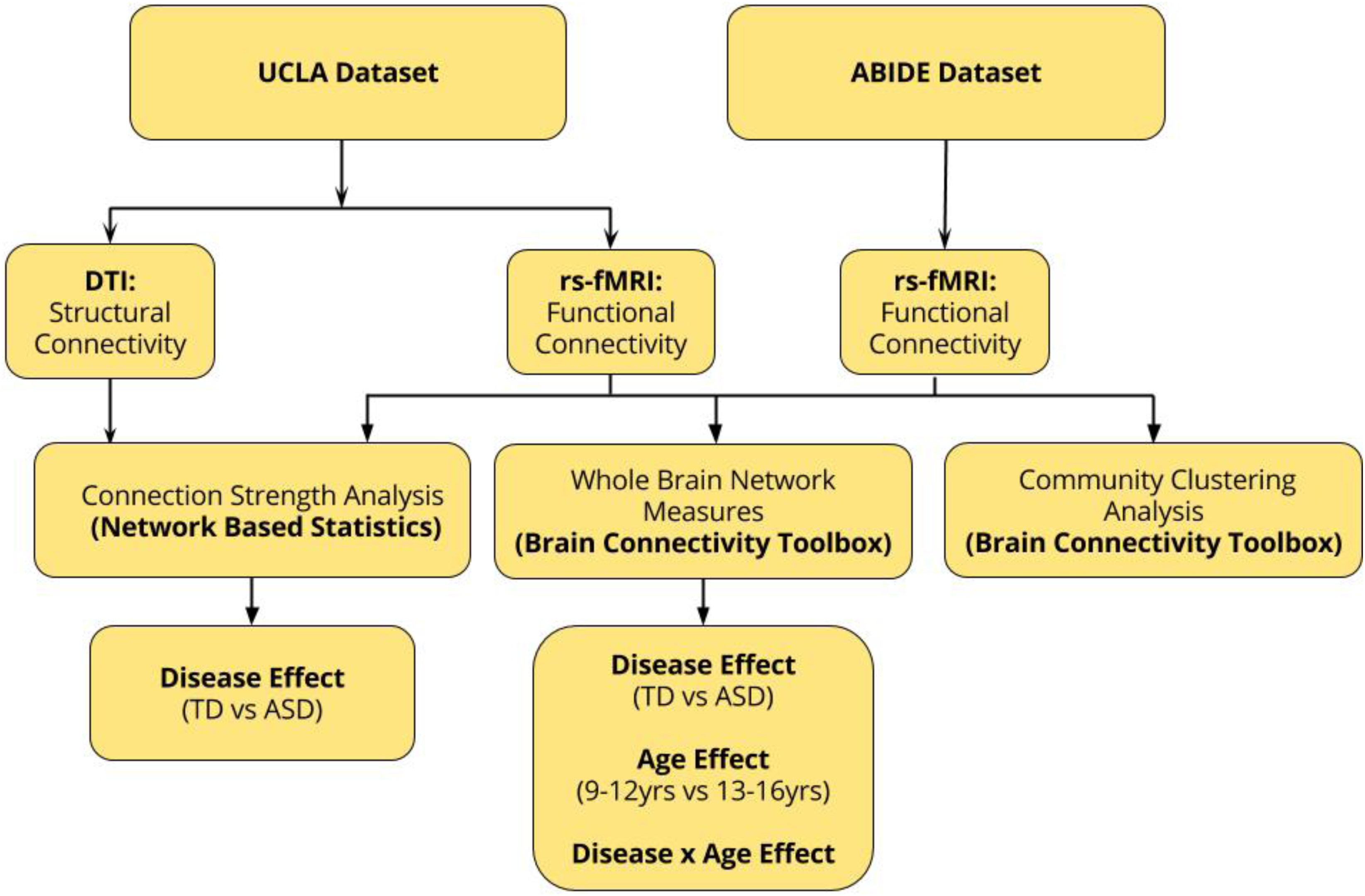
Workflow diagram indicating various analyses carried out on UCLA and ABIDE datasets.

### Clustering Analysis

To construct the most representative structure of network of communities, we applied Louvain algorithm (Blondel et al., 2009) to the unthresholded functional connectivity matrix averaged across all subjects of TD in UCLA and ABIDE datasets. We chose a random partition containing 4 modules (the most common number of modules identified over 100 independent realizations of the algorithm) as our representative partition and reorganized the order of nodes in the FC matrices by this modular organization for visualization purposes. The similarity of this chosen modularity partition with 99 other modularity iterations of the group average matrix was calculated using normalized mutual information and this was used to get the final consensus partition using a threshold of at least 80 similar assignments of an area to an individual cluster over the N=10^3^ runs of the algorithm. For clustering, first an affinity matrix is constructed using radial basis function (RBF)-kernel of the Euclidean distance.

Clustering is done in an algorithmic fashion as listed below:

- Compute un-normalized Laplacian *L*
- Compute first *k* eigenvectors of *L*. Let *U* be a matrix constructed with the above eigenvectors as columns. Cluster the points with K-means clustering algorithm into *K* clusters.

We identified four functional networks and labelled them based on the overlap with consensus mapping provided by Gordon et al. (2014). We observed that the regions assigned to different modules overlap significantly between the Craddock 200 (in ABIDE data set) and Power 264 (in UCLA data set) (**Figure S1**). We found the four predominant communities to be default mode, visual, attention and sensorimotor network. The annotation used for mapping the nodes in ABIDE dataset to functional network is provided in **Supplementary Tables S3**.

### Network-wide group differences

To analyze the edgewise differences in connection strengths and isolate components of the connectivity matrix that differ significantly between ASD and TD, we used permutation testing available in the Network Based Statistic (NBS) Toolbox (Zalesky et al., 2010). The significant differences in each connection value in the functional and structural connectivity matrices of ASD and TD were calculated. Statistics of group differences (TD > ASD & ASD > TD) for edge weights were implemented using one-sided 2-sample t-test, separately for each group using permutation testing. False positives due to multiple comparisons were controlled by using the false discovery rate (FDR corrected for n=10000 iterations, threshold for t-stat=3.0, *p*<0.05), across the whole-brain connectivity matrix. For more stringent thresholding, out of all significant connection differences, only connections above t-stat > 3 are reported. The results are reported for both the UCLA structural and functional data and age-stratified ABIDE functional data. For age stratification, the subjects from ABIDE were divided into four different groups based on disease and age: TD: 9-12 years, TD: 13-16 years, ASD: 9-12 years and ASD: 13-16 years.

### Network Measures

To evaluate the functional networks by means of graph theory, the FC matrices were binarized to adjacency matrices where correlations above a certain threshold are set to 1 and 0 otherwise. For the purpose of our analysis, to define a proper confidence interval, we used 50 thresholds (1%-50%) and reported the average measures across these thresholds. The graph theoretical measures employed were evaluated using Brain Connectivity Toolbox (Rubinov and Sporns, 2010). Binarized FC matrices from each of these groups were then evaluated for six network measures: characteristic path length (CPL), small world index, global efficiency, local efficiency, clustering coefficient and modularity. Characteristic path length (CPL) is the average shortest path length (number of links) between any two nodes in the network. The mean clustering coefficient indicates the probability of two regions that are connected to a third one, also being connected to each other (forming triangles). Mean local efficiency was calculated as the average of inverse shortest path lengths between every pair of nodes which are in the neighborhood of each node. Global efficiency was calculated as the average of inverse shortest path lengths between each pairs of network nodes. The ratio of clustering coefficient and CPL provided the measure for small worldness. Small-worldness evaluates if high global efficiency coexists with high clustering when compared to a random graph. Further, network modularity estimates were computed using the community detection algorithm by Louvain [9]. We have used CPL and global efficiency as measures of functional integration as they estimate the ease with which brain areas can communicate with each other in terms of the path length. Measures of segregation typically quantify the presence of strong densely connected sub-networks within a network. In the context of the brain, this would define the ability to specialize processing within specific modules. Local efficiency, clustering coefficient and modularity are used as measures of functional segregation (Bullmore and Sporns, 2009; Rubinov and Sporns, 2010) (**Table S5)**. These measures were calculated for functional data from both UCLA and age stratified ABIDE datasets.

### Disease and Age-effect on the Network Measures using ABIDE dataset

We used the Stats model python library to perform 2×2 ANOVA to test the effect of disease, age and their interaction on the network integration and segregation measures over the whole brain that are described above. The network measures were calculated at thresholds between 1%-50% and ANOVA test was independently performed at each threshold to study the effect of threshold on the results. Further, to characterize effect of disease, age and their interaction on the local node dynamics, we performed a 2×2 ANOVA on the local efficiency of each node within the identified functional networks.

## Results

### Network-wide group difference analysis

#### UCLA Structural Connectivity

At first, we analyzed UCLA dataset to check whether the structural connectivity between different regions shows similarity and difference across ASD and TD groups. For this purpose, we used t-statistic with FDR correction (*p*<0.05), at different permutation iterations. In structural connectivity matrices, we found significant white matter differences manifested as hypo-connectivity (TD>ASD) between occipital-default mode regions and thalamo-cortical regions. Details of areas can be found in **Table 3**. Similarly, significant hyper-connectivity was found between right putamen and right superior parietal lobule (t-statistic: 3.56, *p*<0.05) (**see Table 3**), which may importantly constrain functions involving spatial attention and numerical ability as observed in Autistic children. Structural hypo-and hyper-connectivity is further displayed in **Figure 2A, B** with a ball and stick diagram by estimating the edge differences between the connected brain regions. Interestingly the atypical connectivity patterns specifically between the subcortico-cortical regions emphasize the importance of these in normative processing.

**Table 3:** Comparison based on t-statistics for the structural connectivity difference in TD > ASD (plain text) and TD <ASD (highlighted) with FDR corrected (p < 0.05) at multiple cluster iterations using UCLA data set.

| Region | Region | t-value |
| --- | --- | --- |
| Left Hippocampus | Right Superior Occipital | 3.41** |
| Left Posterior Cingulate | Right Superior Occipital | 3.51** |
| Right Thalamus | Right Posterior Para hippocampal | 4.61 |
| Right Precentral Cortex | Right Superior Frontal Gyrus | 3.61 |
| Right frontal pole | Left anterior cingulate Cortex | 3.22 |
| **Right Putamen** | **Right Superior Parietal Lobule** | **3.56\*** |

\*\* Survived FDR, 10000 iterations
\* Survived FDR, 8000 iterations

**Fig. 2:**
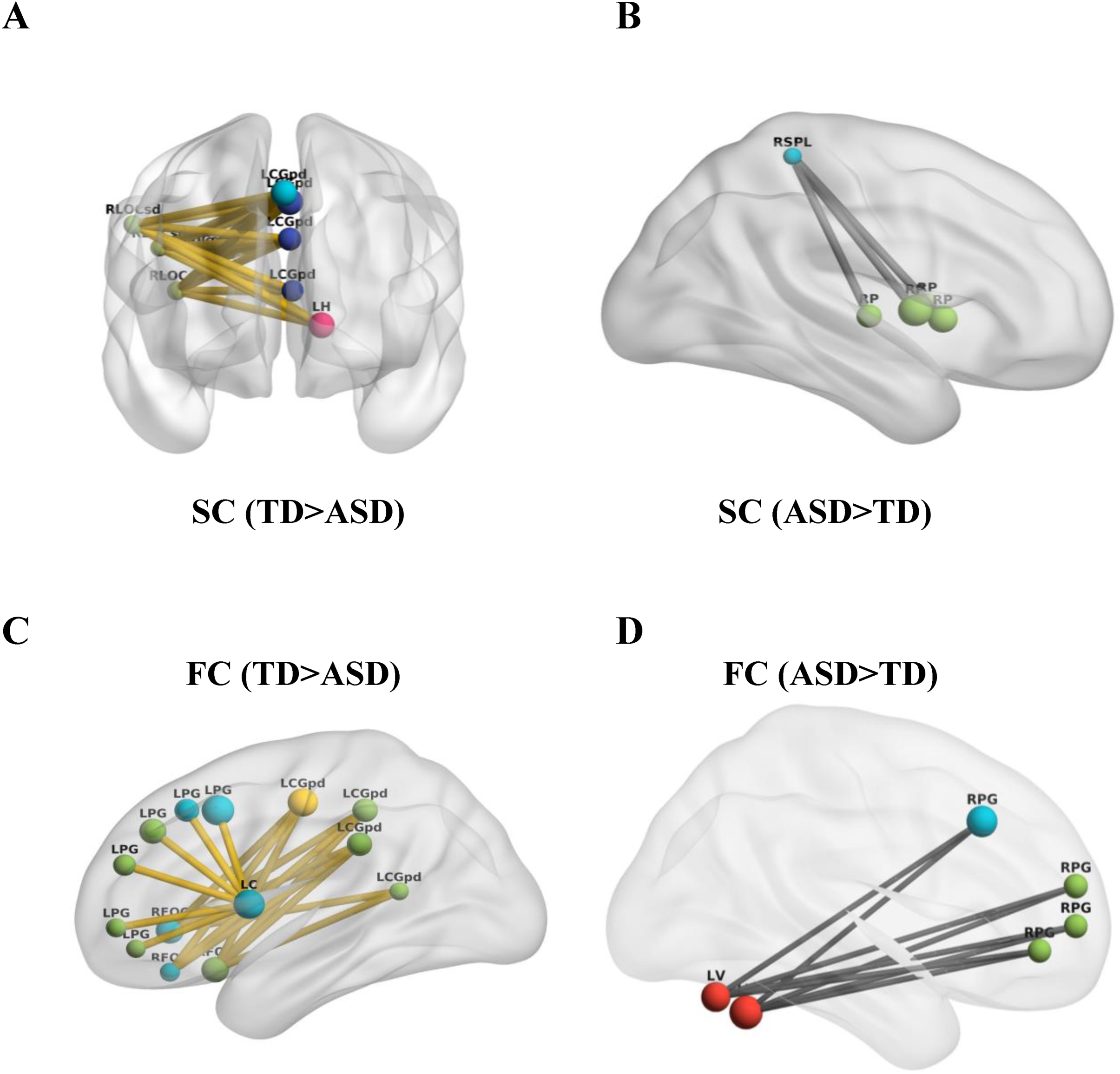
Comparison of structural and functional connectivity edge differences in TD and ASD (UCLA data set). **(A)** TD>ASD anatomical connectivity edge differences are plotted between Default Mode Network, Hippocampus and lateralized Superior Occipital Cortex. (**B)** ASD>TD anatomical connectivity between right Superior Parietal Lobule and Right Putamen indicating abnormal connectivity patterns specially in the proximity of white matter regions of interest. **(C)** TD>ASD functional connectivity edge differences are plotted between Right Frontal Cortex, Left Caudate and Left Paracingulate Gyrus. **(D)** ASD>TD functional connectivity differences are plotted between Right Paracingulate Gyrus and Left Caudate. LCGpd: Left Posterior Cingulate; LH: Left Hippocampus; RLOCsd: Right Lateral Superior Occipital Cortex; LPG: Left Paracingulate Gyrus; LC: Left Caudate; RFOG: Right Frontal Orbital Cortex; RSPL: Right Superior Parietal Lobule; RP: Right Putamen.

#### UCLA Functional Connectivity

Next, we look at the functional connectivity difference between the two groups in order to compare with a number of previous studies. Here, using t-statistic with FDR correction (*p* <0.05), we checked whether the functional connectivity between different regions is significantly similar or different across disease and control groups at various permutation iterations. We found significant functional hypo-connectivity (TD>ASD) between inter-hemispheric default mode connections, for example, between left posterior Cingulate cortex and right precuneus as well as right orbitofrontal cortex, and between left Paracingulate gyrus and left caudate (*p* <0.05). We also found hypo-connectivity between right pre-and post-central gyrus and between left fusiform gyrus and intracalcerine (*p* <0.05). Further details of those areas and results of test statistic for (TD>ASD) can be found in **Table 4 (**and **Figure 2C, D)**. Similarly, significant hyper-connectivity (ASD>TD) was found between many other regions (**Table 4)**. Dysfunction of inter-hemispheric connectivity between specific brain areas may hinder functional integration and segregation and behaviors such as social communication could be compromised. Pairs of region which exhibited structural connectivity differences have also manifested functional connectivity differences which could indicate a correlation between the structure and the function.

**Table 4:** Comparison based on t-statistics for the functional connectivity difference in TD > ASD (plain text) and TD <ASD (highlighted) with FDR corrected (p < 0.05) at multiple cluster iterations using UCLA data set.

| Region | Region | t-value |
| --- | --- | --- |
| Right Frontal Orbital Cortex | Left Posterior Cingulate Gyrus | 3.90* |
| Left Caudate | Left Paracingulate Gyrus | 3.85* |
| Right precuneus | Left posterior cingulate Cortex | 3.70 |
| Right Precentral Cortex | Right Postcentral Cortex | 3.89 |
| Left occipital fusiform gyrus | Left intracalcerine | 3.09 |
| **Right Paracingulate** | **Left Lobule VI of Cerebellum** | **3.94\*** |
| **Left Precuneus** | **Left Frontal Pole** | **3.58** |
| **Right Middle Temporal-occipital** | **Right Anterior Superior Temporal** | **3.05** |
| **Right Putamen** | **Right Superior Parietal Lobule** | **3.14** |

\* Survived FDR, 8000 iterations

#### ABIDE Functional connectivity

Early functional segregation and late functional integration with respect to maturation or development has been shown (Rudie & Dapretto, 2013). Therefore, we studied how disease affects the developmental progression. Using t-statistic with FDR correction (*p* <0.05), across 10000 iterations, we found that for age group 9-12 years there were no connections with ASD>TD or TD>ASD. However, for the adolescent group (13-16 years), there was a significant component (p = 0.042) that survived 10000 iterations having 65 TD>ASD connections identified (**Figure 3A**). Out of these, 24 connections were inter-modular while 41 connections were intra-modular. We further characterized the intra-modular connections within the 4 identified functional networks. Out of the 41 connections, 1 belonged to visual network, 5 to attention and 35 belonged to Default mode network (**Figure 3B**). Further, there were no connections identified with ASD>TD in the adolescent group. These whole brain analyses, albeit restricted to only four resting state networks, indicated the existence of hypo-connectivity in ASD in each of the networks, especially DMN.

**Fig. 3:**
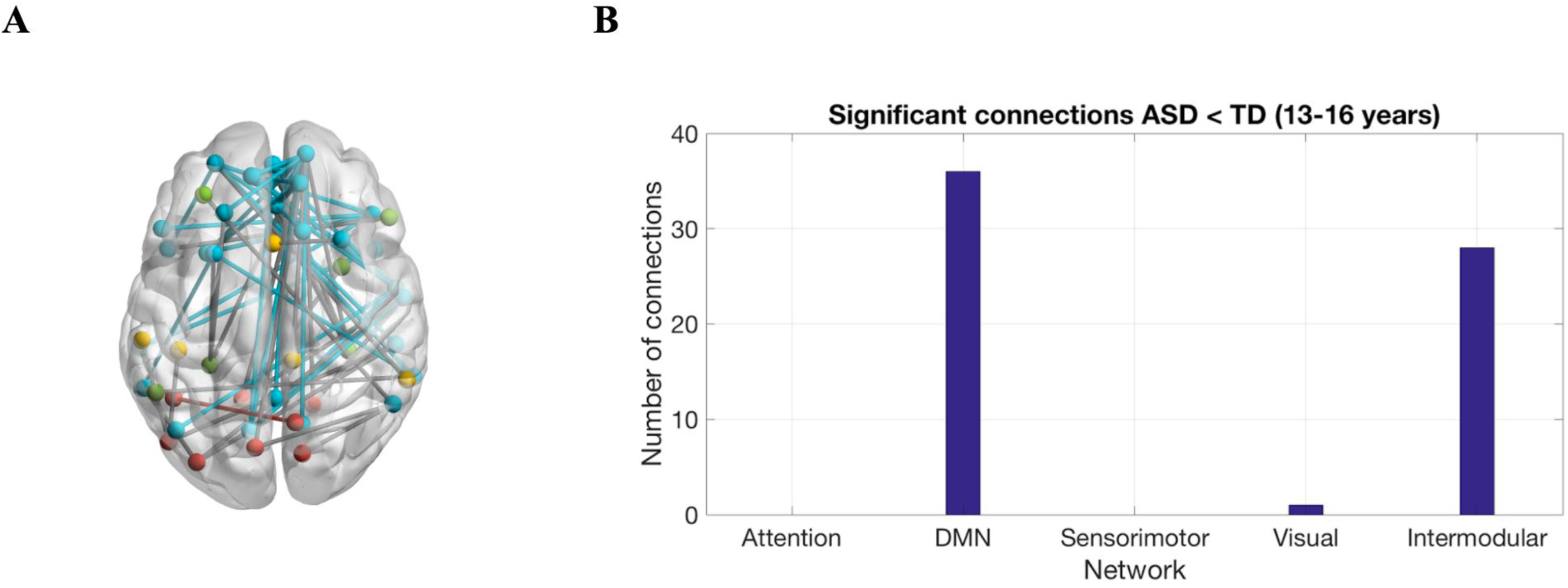
Comparison of functional connectivity edge differences in TD and ASD (ABIDE data set). (A) TD > ASD functional differences in DMN, sensorimotor, attention and visual networks (B) Distribution of TD > ASD within and between functional networks.

### Network measures analysis

The functional integration and segregation can be characterized by various local and global complex network measures (**Table S4**). Therefore, we characterized the differences in the whole brain network between ASD and TD using graph theoretic analysis. In UCLA dataset, we observed significant differences in local (p=0.0044) and global (p=0.0279) efficiency, clustering coefficient (p=0.0074) and modularity (p=0.0058) while CPL and small worldness were not found to be significant (**Table 5 and Figure S2**). In the ABIDE dataset, we observed significant differences in both measures of integration: characteristic path length (CPL, p = 0.03), and measures of segregation: local efficiency (p = 0.02) and modularity (p = 0.02), between TD and ASD only in the adolescent age group (13-16 years) (**Tables 6, 7** and **Figure 4**). In ASD, CPL increased while modularity and local efficiency decreased. On the other hand, small world index and global efficiency did not show significant differences between TD and ASD (**Tables 6, 7)**. The significant differences observed in the graph theoretical measures in adolescents but not in younger children could arise due to significant increase in hypo-connectivity in the four major functional networks. This overall increased hypo-connectivity across networks, may be the cause for abnormally decreased segregation (modularity, local efficiency) and increased integration (CPL) observed in the ASD group (13-16 years) as compared to TD group (13-16 years).

**Table 5:** Network measures for ASD and TD subjects using UCLA data set. *indicates statistical significance.

| **Network Measure** | **TD** | **ASD** | **p-value** |
| --- | --- | --- | --- |
| **Global Efficiency** | 0.5634 | 0.5676 | 0.0279* |
| **Local Efficiency** | 0.7528 | 0.7445 | 0.0044* |
| **CPL** | 1.0444 | 1.2624 | 0.0537* |
| **Modularity** | 0.4105 | 0.3939 | 0.0058* |
| **CC** | 0.5612 | 0.5445 | 0.0074* |
| **Small world index** | 1.8757 | 2.3298 | 0.3346 |

**Table 6:** Network measures for subjects in the age group of 9-12 years using ABIDE data set.

| **Network Measure** | **TD** | **ASD** | **p-value** |
| --- | --- | --- | --- |
| **Global Efficiency** | 0.55794 | 0.558208 | 0.1 |
| **Local Efficiency** | 0.7401 | 0.7382 | 0.21 |
| **CPL** | 0.55092 | 0.54679 | 0.82 |
| **Modularity** | 0.3265 | 0.3200 | 0.12 |
| **CC** | 0.5626 | 0.5611 | 0.74 |
| **Small world index** | 2.7189 | 2.6729 | 0.89 |

**Table 7:**
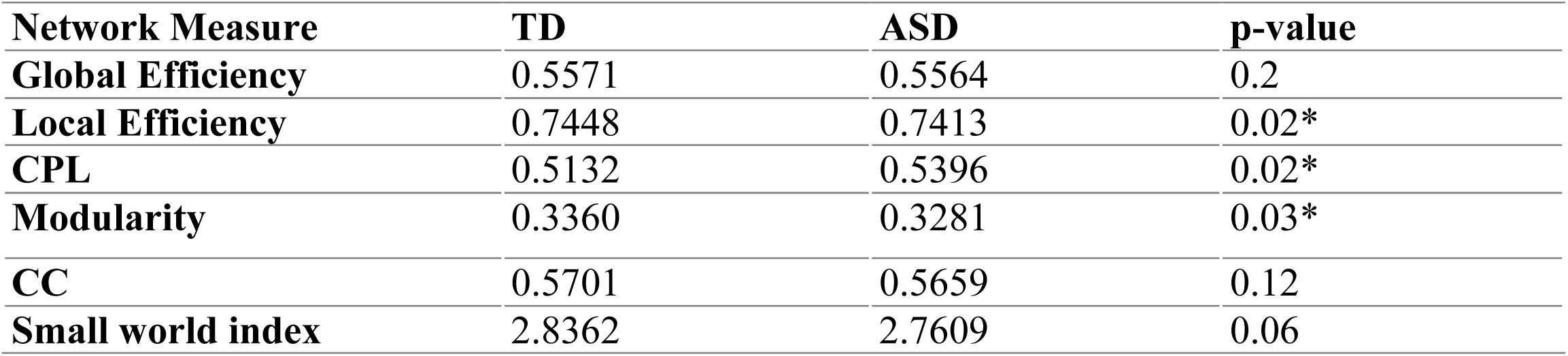
Network measures for subjects in the age group of 13-16 years using ABIDE data set. * indicates statistical significance

| **Network Measure** | **TD** | **ASD** | **p-value** |
| --- | --- | --- | --- |
| **Global Efficiency** | 0.5571 | 0.5564 | 0.2 |
| **Local Efficiency** | 0.7448 | 0.7413 | 0.02* |
| **CPL** | 0.5132 | 0.5396 | 0.02* |
| **Modularity** | 0.3360 | 0.3281 | 0.03* |
| **CC** | 0.5701 | 0.5659 | 0.12 |
| **Small world index** | 2.8362 | 2.7609 | 0.06 |

**Fig. 4:**
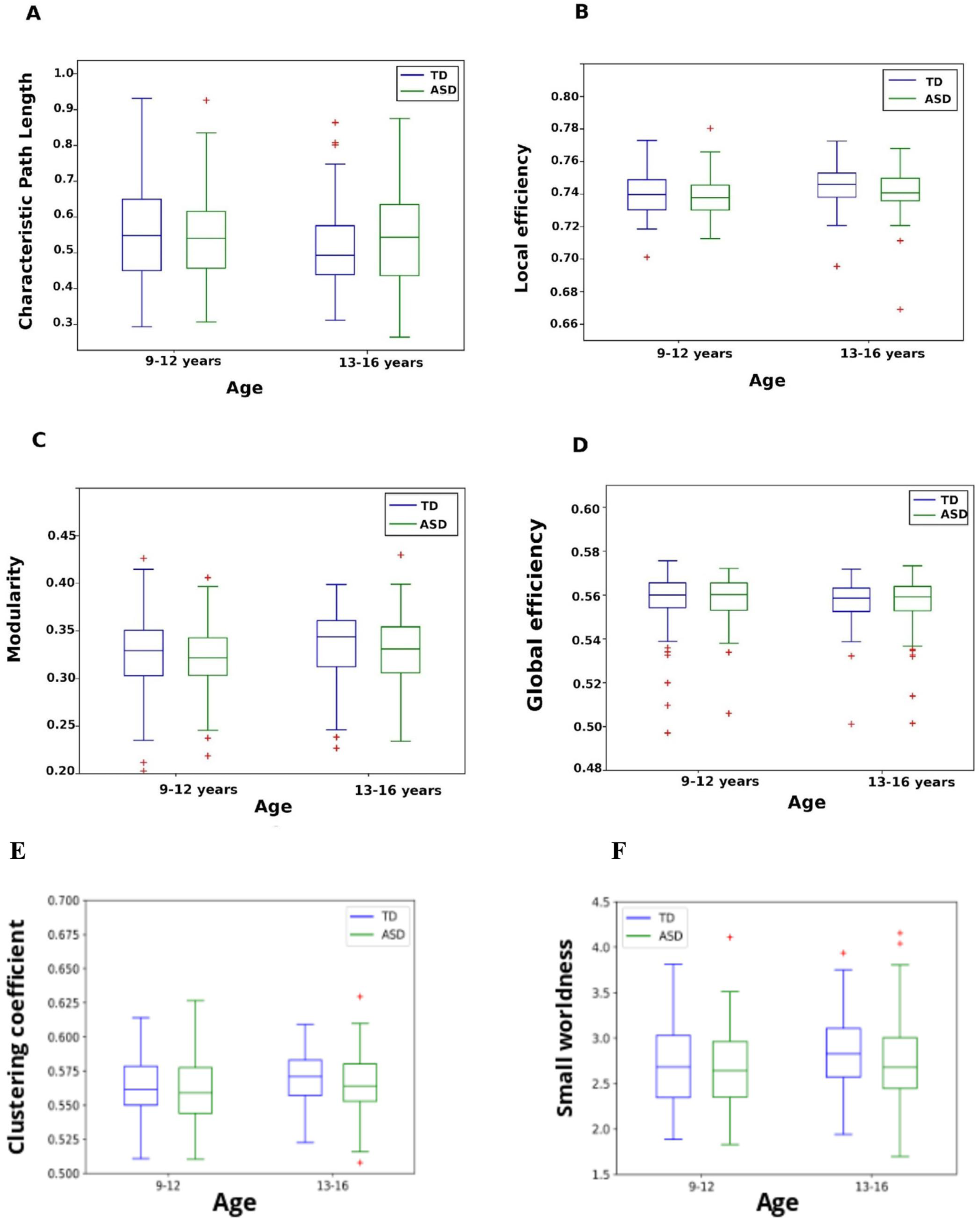
Differences in the network measures of functional integration and segregation between TD and ASD as a function of age. **(A)** Characteristic path length, **(B)** Local efficiency, **(C)** Modularity **(D)** Global efficiency **(E)** Clustering coefficient and **(F)** small worldness (ABIDE data set).

#### Disease, Age, and interaction effects on the Network Measures using ABIDE dataset

2×2 ANOVA revealed that there is significant effect (*p* <0.05) of age and disease on modularity, clustering coefficient, and local efficiency but no interaction effect on these measures (**Figure 5 B, C and D**). CPL and small world index showed significant age effect (**Figure 5A, E**). It can be observed from **Figure 4** that the modularity, local efficiency, clustering coefficient and small world index increased in 13-16 years compared to 9-12 years while CPL decreased in TD. However, the extent of increase (modularity and local efficiency) and decrease (CPL) in these network measures with age is affected in ASD bringing about significant differences between TD and ASD. These differences in the network level measures reflect reduced/delayed functional integration and increased/earlier functional segregation in ASD compared to TD. Further, we observed that these effects on network measures were dependent on the threshold (connection strength) at which they are being tested **(Figure 5)**. The effect of age (threshold=30, F(1,362) =4.3, *p* <0.05) or disease (threshold=30, F(1,362)=4.1, *p* <0.05) on the modularity were observed across higher thresholds (**Figure 5D)**. Further, the effect of age (threshold = 10, F(1,362) = 12.3, *p* <0.05) or disease (threshold=10, F(1,362)=6.5, *p* <0.05) on the local efficiency were observed across lower thresholds (**Figure 5C**). On the other hand, small world index only showed the effect of age (threshold=10, F(1,362)=7.5, *p* <0.05) but not disease across lower thresholds (**Figure 5E**). **Figure S3** shows the sensitivity of these network measures to the range of thresholds. CPL increases as a function of threshold with 13-16 years ASD group exhibiting difference with TD group at higher thresholds (**Figure S3B**). On the other hand, modularity decreases as a function of threshold with 13-16 years ASD group exhibiting higher modularity and less steep asymptote compared to TD (**Figure S3F**).

**Fig. 5:**
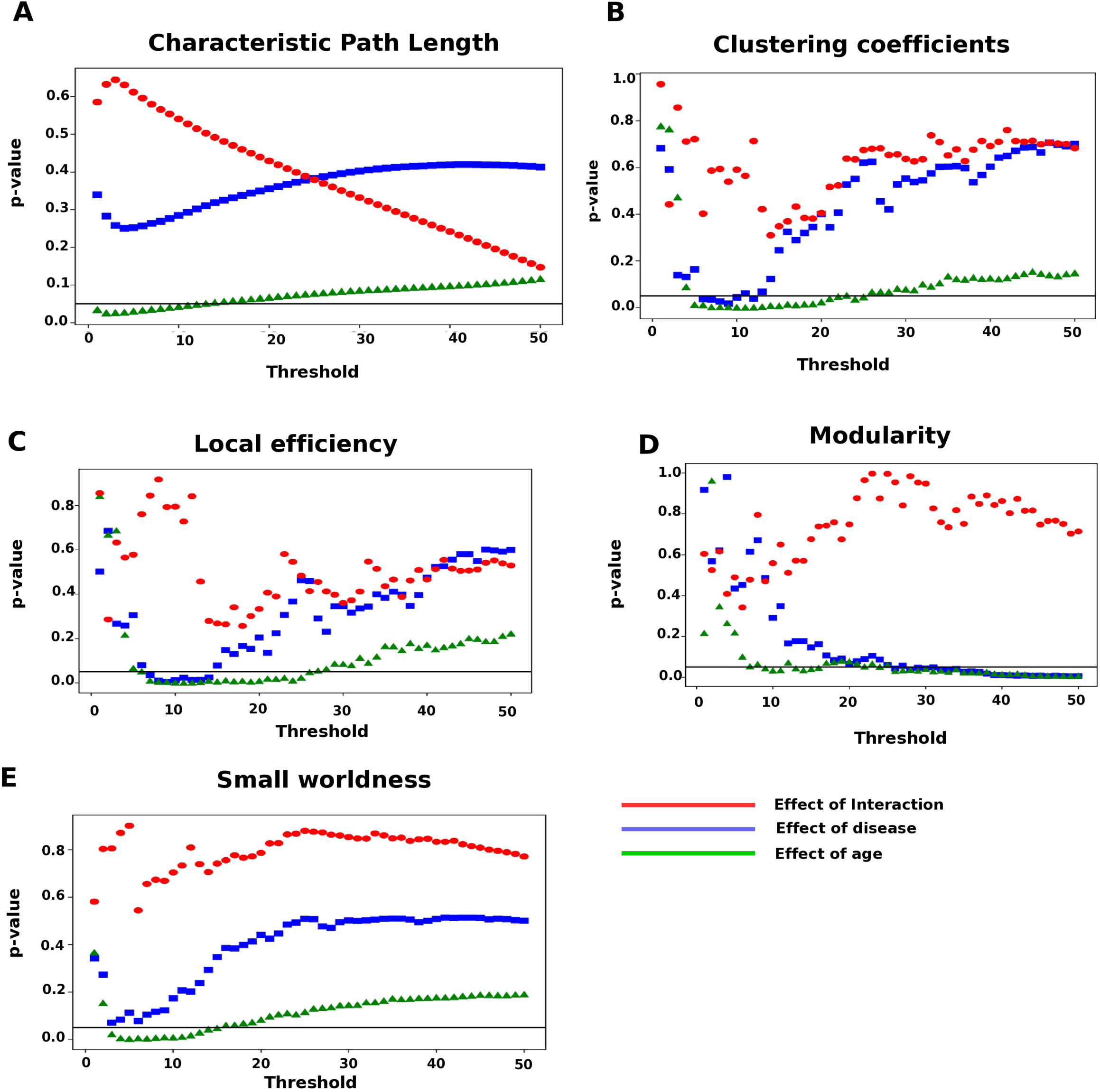
Results of 2×2 ANOVA depicting the effect of age, disease and their interaction on network measures of integration and segregation (ABIDE data set). p-values from ANOVA are plotted for independent life span factors such as disease, age and their interaction as a function of different thresholds for the network measure **(A)** Characteristic path length, **(B)** Clustering coefficients, **(C)** Local efficiency, **(D)** Modularity and **(E)** Small worldness.

Furthermore, we characterized the independent effect of age and disease on functional segregation by performing 2×2 ANOVA to compare local efficiency of each node across age and disease groups. We found that 13 areas belonging to attention (2 areas) and default mode (11 areas) show significant effect of age in the developmental trajectory (9-12 years and 13-16 years). On the other hand, 21 areas belonging to attention (3 areas), default mode (13 areas) and sensorimotor (5 areas) show significant effect of disease in the developmental trajectory (9-12 years and 13-16 years). These regions are shown respectively in **Figure 6A and B**. The combined effect of age and disease was observed in 10 areas belonging to default mode (6 areas), visual (2 areas) and sensorimotor (2 areas) show significant effect of interaction between age and disease (**Figure 6C)**. All these regions are included in **Table 8**.

**Table 8:** Regions of interest (ROIs) in all the four communities (default, attention, visual and sensorimotor) displaying significant age, disease and their interaction effects on local efficiency network measure with 2×2 ANOVA (*p*<0.05).

| Effect of Age | Effect of Disease | Effect of Interaction |
| --- | --- | --- |
| Right Precuneous Cortex | Cingulum_Ant_L | Cingulum_Ant_L |
| Right postcentral gyrus | Left postcentral gyrus | Left middle temporal gyrus |
| Right middle temporal gyrus;<br>posterior division | Cingulum_Ant_R | Frontal_inf_orl_L |
| Left middle frontal gyrus | Right middle frontal gyrus | Left middle cingulate cortex |
| Right inferior frontal gyrus (p.<br>Orbitalis) | Frontal_mid_L | Left superior medial gyrus |
| Left frontal orbital cortex | Right fusiform gyrus | Temporal_mid_L |
| Frontal_inf_Orb_L | Frontal_inf_orb_L | Occipital_Mid_R |
| Left Lateral Occipital Cortex;<br>superior division | Left superior medial gyrus | Left middle occipital gyrus |
| Left superior medial gyrus | Temporal_mid_L | Right precentral gyrus |
| Temporal_mid_L | Left frontal pole | Right superior frontal gyrus |
| Temporal_inf_L | Right middle frontal gyrus |  |
| Left frontal pole | Left frontal orbital cortex |  |
| Temporal_mid_R | Supramarginal_R |  |
|  | Left frontal pole |  |
|  | Temporal_Mid_R |  |
|  | Left inferior frontal gyrus (p.<br>Triangularis) |  |
|  | Right inferior frontal gyrus |  |
|  | Postcentral_L |  |
|  | Right precentral gyrus |  |
|  | Right lateral occipital cortex;<br>superior division |  |
|  | Supp_motor_area_L |  |

**Fig. 6:**
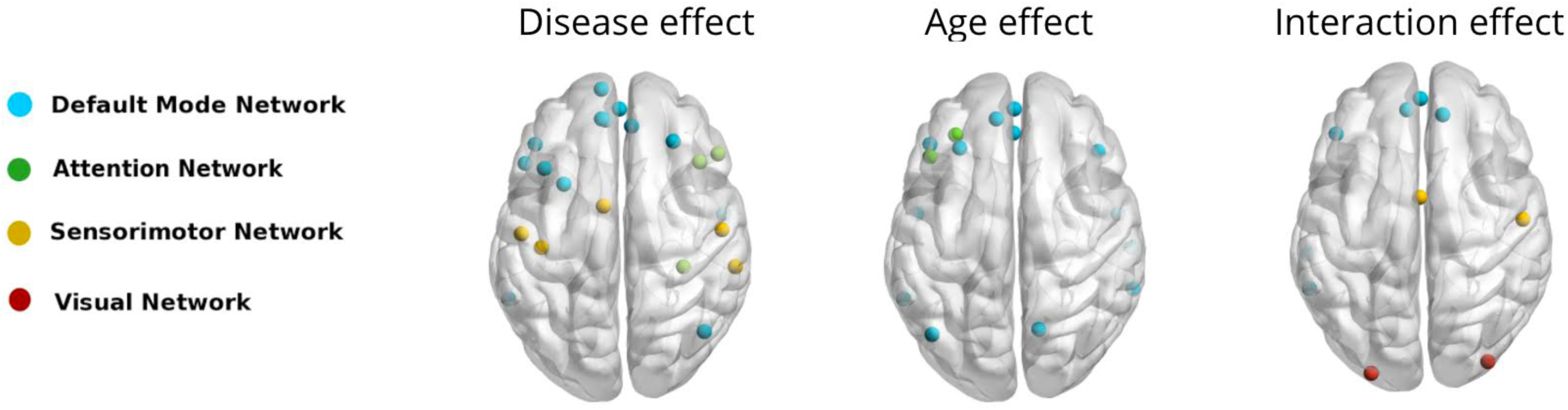
Age, disease and their interaction effects on the local efficiency network measure over the whole brain (ABIDE data set). **(A)** 10 areas of the default network and 10 areas of the attention network show significant effect of age but not disease in the developmental trajectory (9-12 years and 13-16 years). **(B)** 23 areas in total belonging to attention (10 areas), default (9 areas) and sensorimotor (4 areas) show significant effect of disease but not age in the developmental trajectory (9-12 years and 13-16 years). **(C)** 12 areas in total belonging to attention (4 areas), default (4 areas), visual (2 areas) and sensorimotor (2 areas) show significant effect of interaction between age and disease.

## Discussion

Previous studies on whole brain connectivity using UCLA and ABIDE datasets in TD and ASD have shown both under- and over-connectivity (Rudie et al., 2012; Di Martino et al., 2014). Remarkably, many of these earlier reports also found an anterior-posterior gradient of under- to over-connectivity in ASD (Keown et al., 2013; Supekar et al., 2013). A more recent study by Keown et al. (2017) suggests that neither under- nor over-connectivity models fully accommodate the highly diverse evidence from the ASD functional connectivity literature. They found by looking at the graph theoretical properties such as network cohesion and dispersion in the whole brain (distributed over 264 regions of interest comprising 11 resting state networks) that functional networks were globally atypical in ASD, with reduced cohesion and increased dispersion and that connectivity between rich club nodes was atypically increased. In our study, we verify and expand on previous findings by examining the UCLA and ABIDE datasets. By studying the differences in connection strength and network level properties between TD and ASD we found atypical connectivity, both hyper-and hypo-connectivity in ASD whole brain structural and functional networks. We observed that some of the alterations are related to aberrations in structural connectivity. We also found significant effect of age and disease on local and global network measures that characterize the functional integration and segregation. We discuss the key findings of our study below.

### Structural connectivity alterations in ASD

Studies had reported under-connectivity in structural connectivity (Rudie et al., 2012, Supekar et al., 2013, De Martino et al., 2013). Similarly, we also found structural under-connectivity between DMN and occipital regions. Most significant difference was found in connections of the right superior occipital and the (left) default mode regions where Salmi et al., (2013) had previously found group-differences in functional activations. We found evidence for thalamocortical hypo-connectivity in autism as in Nair et al. (2013). The (right) posterior parahippocampal gyrus was found to be under-connected with the (right) thalamus. The parahippocampal gyrus is involved in the functioning of scene recognition (Epstein et al., 1998) and identifying social context as well, including paralinguistic elements of verbal communication such as people’s ability to detect sarcasm. We also found hypo-connectivity between posterior cingulate cortex (PCC) and right occipital cortex, and between right frontal pole and left anterior cingulate cortex. Posterior cingulate cortex (PCC) is the central node in the default mode network (DMN), and its hypo-connectivity is a major bio-marker in detecting ASD. The frontal pole is one of the least understood areas of the brain and the area underwent significant expansion during human evolution. Hypo-connectivity in these regions might point towards abnormal social behavior in these participants. Significant hyper-connectivity between left putamen and left superior parietal lobule might point towards intact motor learning. Interestingly, Travers *et al* also found functional activation in ASD participants in connection with motor learning and repetitive behaviors (Travers et al., 2015) (**Table 3**).

### Functional connectivity alterations in ASD

In ASD, we found significant atypical connectivity, presence of both hyper-and hypo-connectivity in UCLA (**Figure 2C-D, Table 4**) but only hypo-connectivity in ABIDE data sets (**Figure 3)**. This is consistent with different neuroimaging studies that have found increased short-range connections in neurotypical children versus adults (Fair et al., 2009; Supekar et al., 2009; Supekar et al., 2013) but in ASD there are contradictions (Paakki et al., 2010; Shukla et al., 2010; Di Martino et al., 2014). Supekar et al. (2013) showed children with ASD show functional hyper-connectivity whereas Di Martino et al. (2014) reported widespread reductions in connectivity across multiple systems (except for increased connectivity between primary sensory and subcortical regions). Further, two other recent studies (Anderson et al., 2011b; Keown et al., 2013) in ASD also reported widespread reductions in connectivity at both short and long distances using whole-brain approaches to characterize intrinsic connectivity.

We found specifically that the ASD group showed weaker connectivity than TD between the (right) precuneus and the (left) posterior cingulate cortex (PCC) (**Table 4**). In addition, the ASD group showed stronger connectivity relative to the TD between the (left) precuneus and (left) frontal pole. Although some of these regions overlap with structural changes between these regions, most of the other alterations in functional connectivity were not directly related to alterations in fiber density distribution of short and long-range connections. It is interesting that the frontal pole exhibits hyper-connectivity since this has been implicated in the processing of joint attention (Williams et al. (2005)) and found lacking in individuals with ASD.

We observed more significant hypo-connectivity in DMN network in the ABIDE dataset than UCLA (**Fig. 3A versus Fig. 2C**). The nodes that are consistently affected include precuneus, superior frontal, middle temporal, and frontal pole. This analysis suggests that although DMN network is affected in both datasets, the pattern observed is unique and can vary in each Autistic individual. Further, using ABIDE dataset, we found that only the adolescent group (13-16 years), have significant TD>ASD connections. Out of these, 24 connections were inter-modular connections between ROIs belonging to DMN, Attention, Sensorimotor and Visual communities.

### Comparison of network measures of segregation and integration

In examining the differences in six network measures of functional segregation and integration (**Table S4**) between TD and ASD, we observed that in ASD functional segregation decreases while functional integration increases both in UCLA and ABIDE datasets (**Tables 5, 6 and 7**). We found significant differences between TD and ASD in the network measures of segregation such as local efficiency, clustering coefficient and modularity, and in the network measures of integration such as global efficiency (in UCLA) and CPL (in ABIDE). A significant decrease in modularity in ASD suggest a less robust modular organization (i.e. communities were less distinct) in ASD. On the other hand, increase in characteristic path length in ASD suggests more random integration and distribution of edges between nodes. Interestingly, compared to children (9-12 years), adolescents (13-16 years) show remarkable differences in these network measures (**Tables 6, 7**). The overlap and differences in the network measures between the UCLA and ABIDE dataset can also be realized using network based statistics results. Our results are consistent with finding that individuals with ASD exhibit lower clustering (i.e. local efficiency), in particular, in the DMN and secondary Visual communities (Rudie et al., 2012; Keown et al., 2013). Further, our key findings also largely overlap with recent studies on ASD (Supekar et al., 2013; Keown et al., 2013; Doyle-Thomas KA et al., 2015; Hahamy et al., 2015; Abbott et al., 2016).

### Age, disease and their interaction effect on functional segregation and integration

Since age is an important covariate, we studied the effect of age, disease and their interaction on different network measures. Further, we also identified the spatial location and distribution of ROIs in the whole brain network affected by age, pathology and their interaction. We found that measures of segregation (modularity and local efficiency) and a measure of integration (CPL) are significantly affected by age and disease **(Figure 4 and 5)**. With age, we observed that network segregation measures (modularity, clustering coefficient and local efficiency) increases, but in ASD the extent of increase was affected. Our findings lend support to a model of global atypical connections during maturation similar to what has been reported recently using whole brain network analysis (Keown et al., 2017).

Further, the results of 2×2 ANOVA indicate that in ASD the local efficiency of individual nodes of DMN, sensorimotor and attention network decreased suggesting a decrease in the efficiency of functional segregation (**Table 8**). The effect of maturation on local efficiency seems to be prominent in the cortical circuits with the frontal pole and nearby frontal regions showing age-related differences. This is in line with the developmental trajectory where the most anterior regions of the frontal lobe mature at the last. The effect of disease seems to be related to both the anterior regions such as the frontal regions as well as the posterior cortical regions such as the precuneus (parietal) and occipital regions. Disease seems to impact, in addition to the superior and inferior frontal regions, regions in the temporal lobe including the boundary region such as fusiform gyrus, parietal region such as supramarginal gyrus and supplementary motor area. The age and disease interaction seems to be related to all the networks such as DMN, sensorimotor, and visual. Overall these areas correlate well with the behavioral traits related to language/communication, sensory and attentional disturbances and repetitive motor actions.

Our study differed in several respects from previous graph theory investigations in ASD. We verified our finding across multiple datasets. We keep a larger confidence interval by considering all thresholds from 1%-50% while computing network measures. However, we do not apply sparsity threshold criterion while calculating group differences that may reduce sensitivity not only to inter-individual differences due to noise, but also to global group (typical vs. atypical) connectivity differences (e.g., predominant under-connectivity or over-connectivity in ASD) (Keown et al., 2017). Further, we go beyond the models proposing general functional under-connectivity or general over-connectivity in ASD as they are probably too simple to account for the complex patterns of connectivity.

### Limitations and Future work

Complex network analysis, being a recent approach, lacks standardization of methods. Whole brain connectivity differences are highly sensitive to algorithms used for thresholding and partitioning graphs into community structures. For better insight and reproducibility of results across studies and age groups, higher-order statistics are required. False discovery rate (FDR) methods used for validation of t-statistic are highly sensitive to the number of permutations. Therefore, results are prone to yielding false positives.

Our work is a step in that direction and we take these results with plenty of caution. To consistently find significant difference in the same graph theoretic network measures of segregation and integration, we validated our results with two independent samples. One needs care about the validation samples as ABIDE consists of data sets collected from multiple sites. We did not study the correlation of the graph measures with the symptom severity (ADOS scores) because ADOS were not available for majority of samples.

Further, there are significant group differences in verbal IQ between subjects. A previous study on UCLA dataset concluded that defects in default and visual system largely co-varied with the social symptom severity and not with the communicative symptom severity (Rudie et al., 2012, 2013). However, whether verbal fluency shows any significant correlation with the identified functional hypo-connectivity and network measures along with age, disease and interaction in the individuals with ASD is still an open question and needs future study.

Another important limitation of this study is that our ASD sample did not represent full spectrum of autistic disorders as the data in the low functioning autistic samples are usually full of motion artefacts. Here, we have tried to hand pick datasets from multiple sites which are pre-processed and minimally affected by motion. In this process, perhaps we have included a large number of high functioning autistic individuals in the sample. However, our samples also include low functioning autistic individuals. Little is known from neuroimaging studies the effect of age and disease in low functioning autistic individuals, and our findings could fill that gap. Importantly, closer methodological scrutiny is obviously required, given the recent controversy regarding the effects of motion confounds, whereby not appropriately correcting for head motion can lead to both spurious increases in local connectivity and reductions in long-range connectivity. In this context, ABIDE used advanced motion-correction techniques and addressed other major methodological concerns (i.e., global signal regression) before the public release of dataset for reproducibility research.

An interesting observation arising from the analysis of rs-fMRI of ASD is that the results point to the existence of both hypo-and hyper-connectivity. An intriguing possibility is that this is due to variations in the functioning-level of the ASD participants, i.e., a group of low functioning individuals could show hypo-FC as against hyper-FC in the case of high functioning ASD subjects. However, since ADOS scores are not available for a majority of samples, this possibility could not be verified. Even though functional connectivity is shaped by the underlying structure, it cannot be fully explained just by structure. One way to examine finer scale differences in the resting state dynamics of the ASD and TD children is therefore to use biologically plausible whole-brain computational models which can be compared across populations or for investigation of individual damage in structural connectivity modules as well as in rehabilitation (Vattikonda et al., 2016).

## Author Contributions

DR, SBR, PKV have designed the research problem. VH, SN, PKV, SBR, DR carried out the original research. VH, SN analyzed data. VH, SN, PKV, SBR, DR have all contributed in writing this original research article.

## Funding

DR is supported by the Ramalingaswami fellowship (BT/RLF/Re-entry/07/2014) from Department of Biotechnology (DBT), Ministry of Science & Technology, Government of India. PKV acknowledges financial support from the Early Career Research Award Scheme, Science and Engineering Research Board, DST, India.

### Acknowledgements

The support of Pramod S. K. and Shailesh Jain of Cognitive Science Lab, IIIT Hyderabad for a partial analysis of the data and generation of figures is gratefully acknowledged.

## Supplementary Tables

**Table S1:** Site specific data for ABIDE (9-12 years). FD: Mean Framewise Displacement, FIQ: Full Scale IQ score, VIQ: Verbal IQ score, PIQ: Performance IQ score

| Sites | Sample size |  | Age<br>mean(SD)<br>[range] |  | FD<br>mean (SD)<br>[range] |  | FIQ<br>mean (SD)<br>[range] |  | VIQ<br>mean (SD)<br>[range] |  | PIQ<br>mean (SD)<br>[range] |  | Handedness |  | Gender |  |
| --- | --- | --- | --- | --- | --- | --- | --- | --- | --- | --- | --- | --- | --- | --- | --- | --- |
|  | ASD | TD | ASD | TD | ASD | TD | ASD | TD | ASD | TD | ASD | TD | Left | Right | Male | Female |
| **Stanford** | 9 | 12 | 10.566<br>(1.3)<br>[8.42-11.8] | 10.1<br>(0.75)<br>[9.24-11.69] | 0.086<br>(0.033)<br>[0.037-0.1119] | 0.09<br>(0.049)<br>[0.03-0.18] | 107.75<br>(20.968)<br>[78-137] | 107.27<br>(16.59)<br>[79-124] | 100.75<br>(24.13)<br>[72-137] | 103.6<br>(19.4)<br>[67-136] | 114.25<br>(15.2)<br>[89-128] | 109.4<br>(18.1)<br>[81-145] | 2 | 14 | 11 | 5 |
| **USM** | 11 | 7 | 10.3<br>(1.82)<br>[9.5 -<br>11.5] | 11.1<br>(1.3)<br>[9.3-11.94] | 0.055<br>(0.04)<br>[0.050-0.058] | 0.166<br>(0.05)<br>[0.03-0.19] | 100.3<br>(12.3)<br>[88-113] | 144<br>(15.3)<br>[120-150] | 97.3<br>(18.3)<br>[79-103] | 120 (13.2)<br>[85-153] | 109.5<br>(10.2)<br>[85-115] | 138<br>(18.6)<br>[110-160] | NA | NA | 10 | 2 |
| **Pitt** | 10 | 7 | 10.6 (1.3)<br>[10.2-11.8] | 11.41<br>(1.2)<br>[9.8-12.8] | 0.133<br>(0.032)<br>[0.07-0.18] | 0.1916<br>(0.027)<br>[0.091-0.15] | 106.5<br>(10.5)<br>[96-117] | 107<br>(10.3)<br>[88-118] | 103.5<br>(15.5)<br>[88-120] | 98 (13.2)<br>[84-110] | 108.5<br>(8.5)<br>[100-117] | 11754<br>(13.21)<br>[90-134] | NA | NA | 14 | 2 |
| **NYU** | 28 | 18 | 10.04<br>(0.955)<br>[8.51-11.92] | 9.972<br>(1.295)<br>[9.01-11.92] | 0.088<br>(0.047)<br>[0.03-0.189] | 0.056<br>(0.024)<br>[0.021-0.12] | 106.7<br>(16.01)<br>[76-148] | 112.92<br>(15.1)<br>[80-142] | 102.7<br>(14.41)<br>[77-137] | 112.71<br>(15.226)<br>[85-143] | 110.3<br>(18.26)<br>[72-149] | 110.32<br>(14.9)<br>72-135] | NA | NA | 27 | 1 |
| **UM** | 10 | 15 | 10.5<br>(0.84)<br>[9.2-11.5] | 10.47<br>(0.93)<br>[9.2-12.8] | 0.11<br>(0.05)<br>[0.02-0.19] | 0.068<br>(0.036)<br>[0.029-0.15] | 103.05<br>(21.7)<br>[78.5-147.5] | 107.333<br>(9.97)<br>[92.5-127.5] | 109.3<br>(27.44)<br>[82-180] | 115<br>(11.83)<br>[102-138] | 96.7<br>(22.7)<br>[64-137] | 99.06<br>(9.73)<br>[82-117] | 2 | 7 | 6 | 4 |
| **Yale** | 20 | 14 | 10.87<br>(1.11)<br>[8.92-11.92] | 10.907<br>(0.93)<br>[9.42-12.5] | 0.0126<br>(0.044)<br>[0.078-0.194] | 0.085<br>(0.047)<br>[0.039-0.19] | 93.28<br>(18.24)<br>[61-121] | 104.222<br>(14.62)<br>[78-127] | 101.2<br>(17.87)<br>[66-126] | 110.3<br>(13.2)<br>[89-1132] | 89.85<br>(14.87)<br>[60-107] |  | 1 | 19 | 17 | 3 |

**Table S2:**
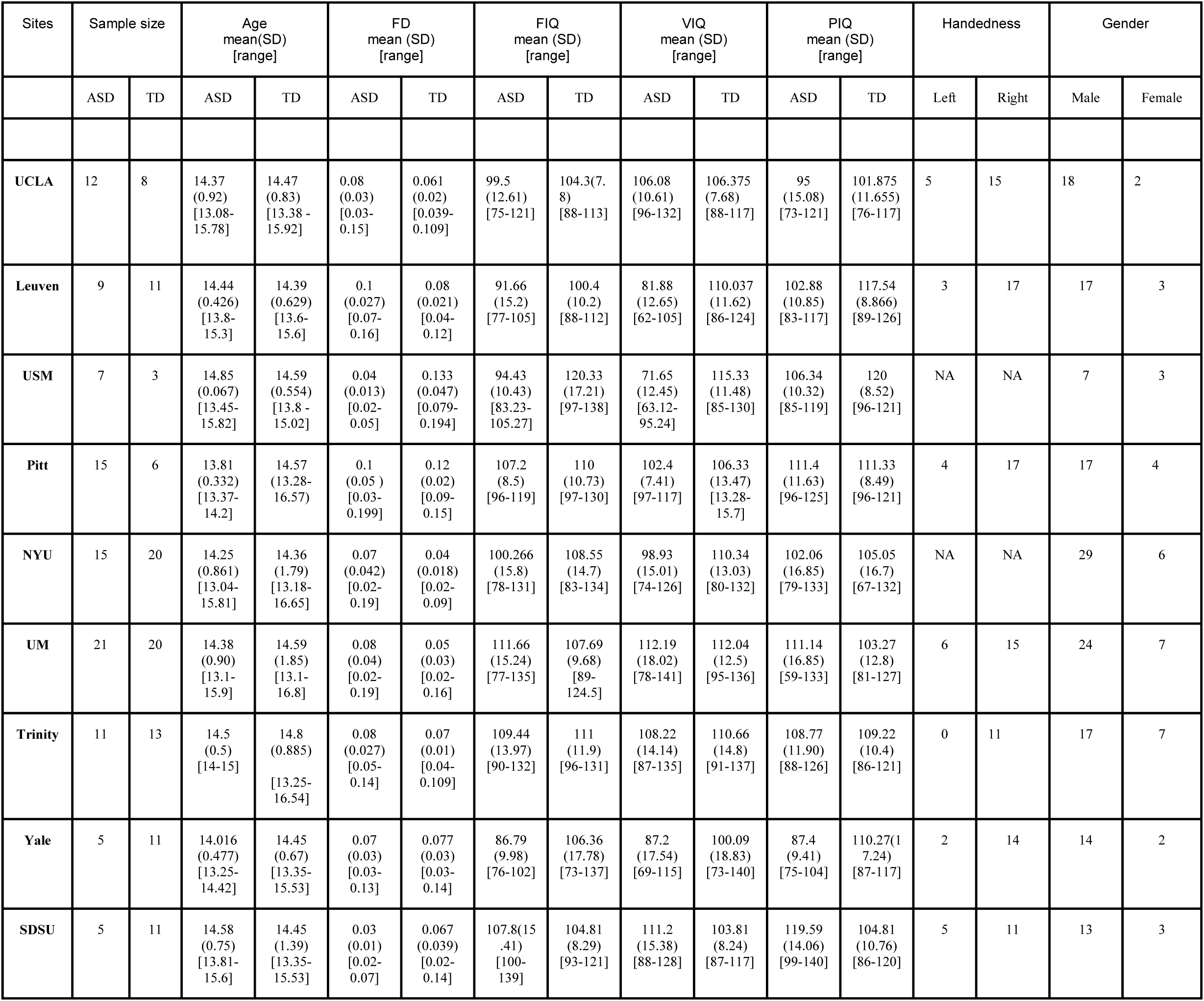
Site Specific Data for ABIDE (13-16 years). FD: Mean Framewise Displacement, FIQ: Full Scale IQ score, VIQ: Verbal IQ score, PIQ: Performance IQ score

**Table S3:** Consensus for ABIDE dataset (Craddock parcellation)

| ROI name | Network |
| --- | --- |
| Right Temporal Occipital Fusiform Cortex | Visual |
| Precentral_L | Sensorimotor |
| Cingulum_Ant_L | Attention |
| Cingulum_Ant_R | Attention |
| Postcentral_L | Sensorimotor |
| Frontal_Inf_Orb_L | DMN |
| Right Frontal Pole | Salience |
| Insula_L | Salience |
| Left Fusiform Gyrus | Visual |
| Left Lingual Gyrus | Visual |
| Right Cingulate Gyrus; posterior division | Unknown |
| Insula_R | Cingular Opercular |
| Right Medial Frontal Gyrus | Attention |
| Temporal_Inf_L | Attention |
| Left Frontal Orbital Cortex | DMN |
| Temporal_Inf_R | Unknown |
| Occipital_Inf_R | Visual |
| Right Superior Temporal Gyrus | Attention |
| Left Thalamus | Subcortical |
| Left SupraMarginal Gyrus | Attention |
| Left Precentral Gyrus | Sensorimotor |
| Parietal_Inf_L | DMN |
| Right Inferior Parietal Lobule | Sensorimotor |
| Left Inferior Frontal Gyrus (p. Opercularis) | Attention |
| Right Subcallosal Cortex | Unknown |
| Right Posterior Cingulate | DMN |
| Right Insula Lobe | Unknown |
| Right Declive | Unknown |
| Cingulum_Mid_L | DMN |
| Right Occipital Pole | Visual |
| Left Superior Temporal Gyrus | DMN |
| Right Lingual Gyrus | Visual |
| Left Cerebellum | Cerebeller |
| Left Middle Occipital Gyrus | Visual |
| Right Middle Temporal Gyrus; posterior division | DMN |
| Left Lateral Occipital Cortex; superior division | Visual |
| RightPrecentral Gyrus | Attention |
| Right Uncus | Unknown |
| Cingulum_Mid_R | DMN |
| Right Subcallosal Gyrus | DMN |
| SupraMarginal_L | Attention |
| Right Precuneous Cortex | DMN |
| SupraMarginal_R | Sensorimotor |
| Right Anterior Cingulate Cortex | DMN |
| Left Culmen | Unknown |
| Right Thalamus | Subcortical |
| Right Inferior Frontal Gyrus | Attention |
| Parietal_Sup_R | Sensorimotor |
| Left Uncus | Unknown? |
| Right Putamen | subcortical |
| Right Temporal Pole | DMN |
| Right Inferior Frontal Gyrus (p. Orbitalis) | DMN |
| Right Lateral Occipital Cortex; superior division | Attention |
| Left Middle Frontal Gyrus | Attention |
| Caudate_R | Subcortical |
| Caudate_L | Subcortical |
| Right Postcentral Gyrus | Sensorimotor |
| Supp_Motor_Area_L | Sensorimotor |
| Right Middle Frontal Gyrus | DMN |
| Right Insula | Salience |
| Right SupraMarginal Gyrus | Attention |
| Supp_Motor_Area_R | Sensorimotor |
| Right Culmen | Unknown |
| Right Middle Temporal Gyrus | DMN |
| Right Inferior Frontal Gyrus (p. Opercularis) | DMN |
| Left Cuneus | DMN |
| Left Temporal Pole | DMN |
| Left Superior Medial Gyrus | DMN |
| Right Parahippocampal Gyrus | DMN |
| Left Inferior Frontal Gyrus (p. Triangularis) | DMN |
| Right Cuneus | DMN |
| Calcarine_L | Visual |
| Left Middle Temporal Gyrus | DMN |
| Left Precuneous Cortex | DMN |
| Left Postcentral Gyrus | Sensorimotor |
| Left Precuneus | DMN |
| Right Paracingulate Gyrus | DMN |
| Right Inferior Frontal Gyrus (p. Triangularis) | Salience |
| Left Inferior Parietal Lobule | DMN |
| Left Superior Parietal Lobule | Attention |
| Left Parahippocampal Gyrus | Unknown |
| Left Superior Frontal Gyrus | DMN |
| Occipital_Mid_R | Visual |
| Right Fusiform Gyrus | Visual |
| Right Superior Frontal Gyrus | DMN |
| Left Frontal Pole | DMN |
| Right Lateral Occipital Cortex; inferior division | Attention |
| Frontal_Mid_L | DMN |
| Left Lateral Occipital Cortex; inferior division | Attention |
| Left Occipital Pole | Visual |
| Left Putamen | Subcortical |
| Left Inferior Frontal Gyrus | DMN |
| Left Inferior Temporal Gyrus | DMN |
| Temporal_Mid_L | DMN |
| Left Middle Cingulate Cortex | DMN |
| Right Caudate Nucleus | Subcortical |
| Temporal_Mid_R | DMN |
| Left Medial Frontal Gyrus | DMN |
| Left Insula | Salience |
| Right Precentral Gyrus | Sensorimotor |
| Right Cingulate Gyrus; anterior division | Sensorimotor |
| Right Frontal Orbital Cortex | Unknown |
| Right Precuneus | DMN |

**Table S4:** Consensus for UCLA dataset (Power parcellation)

| **ROI name** | **Network** |
| --- | --- |
| Left Occipital Pole | Uncertain |
| Right Frontal Orbital Cortex | Uncertain |
| Left Inferior Temporal Gyrus temporooccipital part | Uncertain |
| Right Frontal Medial Cortex | Uncertain |
| Left Parahippocampal Gyrus anterior division | Uncertain |
| Right Parahippocampal Gyrus posterior division | Uncertain |
| Left Temporal Fusiform Cortex posterior division | Uncertain |
| Right Middle Temporal Gyrus posterior division | Uncertain |
| Right Inferior Temporal Gyrus posterior division | Uncertain |
| Right Frontal Pole | Uncertain |
| Left Precuneous Cortex | Sensory/somatomotor Hand |
| Left Precentral Gyrus | Sensory/somatomotor Hand |
| Left Cingulate Gyrus posterior division | Sensory/somatomotor Hand |
| Right Cingulate Gyrus anterior division | Sensory/somatomotor Hand |
| Left Precentral Gyrus | Sensory/somatomotor Hand |
| Left Postcentral Gyrus | Sensory/somatomotor Hand |
| Right Postcentral Gyrus | Sensory/somatomotor Hand |
| Right Precentral Gyrus | Sensory/somatomotor Hand |
| Right Superior Parietal Lobule | Sensory/somatomotor Hand |
| Left Superior Parietal Lobule | Sensory/somatomotor Hand |
| Right Insular Cortex | Sensory/somatomotor Mouth |
| Left Juxtapositional Lobule Cortex | Cingulo-opercular Task Control |
| Right Supramarginal Gyrus anterior division | Cingulo-opercular Task Control |
| Right Superior Frontal Gyrus | Cingulo-opercular Task Control |
| Left Superior Frontal Gyrus | Cingulo-opercular Task Control |
| Right Juxtapositional Lobule Cortex | Cingulo-opercular Task Control |
| Left Central Opercular Cortex | Cingulo-opercular Task Control |
| Right Central Opercular Cortex | Cingulo-opercular Task Control |
| Left Insular Cortex | Cingulo-opercular Task Control |
| Left Cingulate Gyrus anterior division | Cingulo-opercular Task Control |
| Right Insular Cortex | Auditory |
| Right Superior Temporal Gyrus posterior division | Auditory |
| Right Planum Temporale | Auditory |
| Left Planum Temporale | Auditory |
| Right Occipital Pole | Auditory |
| Right Parietal Operculum Cortex | Auditory |
| Left Parietal Operculum Cortex | Auditory |
| Left Central Opercular Cortex | Auditory |
| Right Central Opercular Cortex | Auditory |
| Right Postcentral Gyrus | Auditory |
| Left Insular Cortex | Auditory |
| Left Lateral Occipital Cortex superior division | Default mode |
| Right Frontal Pole | Default mode |
| Right Frontal Medial Cortex | Default mode |
| Left Hippocampus | Default mode |
| Left Frontal Pole | Default mode |
| Left Lateral Occipital Cortex superior division | Default mode |
| Right Lateral Occipital Cortex superior division | Default mode |
| Left Temporal Pole | Default mode |
| Right Temporal Pole | Default mode |
| Left Middle Temporal Gyrus posterior division | Default mode |
| Right Frontal Orbital Cortex | Uncertain |
| Left Lateral Occipital Cortex superior division | Default mode |
| Left Precuneous Cortex | Default mode |
| Right Precuneous Cortex | Default mode |
| Left Cingulate Gyrus posterior division | Default mode |
| Right Cingulate Gyrus posterior division | Default mode |
| Right Precuneous Cortex | Default mode |
| Right Lateral Occipital Cortex superior division | Default mode |
| Right Superior Frontal Gyrus | Default mode |
| Left Superior Frontal Gyrus | Default mode |
| Left Middle Frontal Gyrus | Default mode |
| Right Frontal Pole | Default mode |
| Left Frontal Pole | Default mode |
| Left Paracingulate Gyrus | Default mode |
| Right Paracingulate Gyrus | Default mode |
| Left Cingulate Gyrus anterior division | Default mode |
| Left Frontal Pole | Default mode |
| Right Middle Temporal Gyrus posterior division | Default mode |
| Left Middle Temporal Gyrus posterior division | Default mode |
| Right Cingulate Gyrus anterior division | Default mode |
| Right Superior Temporal Gyrus anterior division | Default mode |
| Left Lingual Gyrus | Default mode |
| Right Parahippocampal Gyrus posterior division | Default mode |
| Left Temporal Fusiform Cortex posterior division | Default mode |
| Right Crus I | Default mode |
| Right Temporal Pole | Default mode |
| Left Middle Temporal Gyrus anterior division | Default mode |
| Right Angular Gyrus | Default mode |
| Left Middle Temporal Gyrus posterior division | Default mode |
| Left Cingulate Gyrus posterior division | Memory retrieval |
| Left Precuneous Cortex | Memory retrieval |
| Right Precuneous Cortex | Memory retrieval |
| Left Frontal Orbital Cortex | Default mode |
| Left Superior Frontal Gyrus | Ventral attention |
| Right Occipital Pole | Uncertain |
| Left Occipital Pole | Uncertain |
| Right Lateral Occipital Cortex inferior division | Visual |
| Right Intracalcarine Cortex | Visual |
| Left Intracalcarine Cortex | Visual |
| Left Lateral Occipital Cortex superior division | Visual |
| Left Occipital Pole | Visual |
| Right Temporal Occipital Fusiform Cortex | Visual |
| Left Lateral Occipital Cortex inferior division | Visual |
| Right Lateral Occipital Cortex superior division | Visual |
| Right Occipital Fusiform Gyrus | Visual |
| Right Cuneal Cortex | Visual |
| Right Lateral Occipital Cortex inferior division | Visual |
| Right Lateral Occipital Cortex superior division | Visual |
| Right Cuneal Cortex | Visual |
| Left Lateral Occipital Cortex inferior division | Visual |
| Right Occipital Fusiform Gyrus | Visual |
| Left Cuneal Cortex | Visual |
| Left Occipital Fusiform Gyrus | Visual |
| Right Lateral Occipital Cortex inferior division | Visual |
| Left Precentral Gyrus | Fronto-parietal Task Control |
| Right Middle Frontal Gyrus | Fronto-parietal Task Control |
| Left Inferior Frontal Gyrus pars opercularis | Fronto-parietal Task Control |
| Left Supramarginal Gyrus posterior division | Fronto-parietal Task Control |
| Left Superior Frontal Gyrus | Fronto-parietal Task Control |
| Right Inferior Temporal Gyrus temporooccipital part | Fronto-parietal Task Control |
| Right Frontal Pole | Fronto-parietal Task Control |
| Left Occipital Fusiform Gyrus | Uncertain |
| Right Crus II | Uncertain |
| Right Crus I | Uncertain |
| Right Precentral Gyrus | Fronto-parietal Task Control |
| Left Middle Frontal Gyrus | Fronto-parietal Task Control |
| Left Frontal Pole | Fronto-parietal Task Control |
| Right Frontal Pole | Fronto-parietal Task Control |
| Right Supramarginal Gyrus posterior division | Fronto-parietal Task Control |
| Left Superior Parietal Lobule | Fronto-parietal Task Control |
| Right Angular Gyrus | Fronto-parietal Task Control |
| Right Superior Parietal Lobule | Fronto-parietal Task Control |
| Left Middle Frontal Gyrus | Fronto-parietal Task Control |
| Right Supramarginal Gyrus posterior division | Salience |
| Right Precentral Gyrus | Salience |
| Right Middle Frontal Gyrus | Salience |
| Right Inferior Frontal Gyrus pars triangularis | Salience |
| Left Insular Cortex | Salience |
| Right Insular Cortex | Salience |
| Right Frontal Orbital Cortex | Salience |
| Right Insular Cortex | Salience |
| Left Cingulate Gyrus anterior division | Salience |
| Left Paracingulate Gyrus | Salience |
| Left Frontal Pole | Salience |
| Left Cingulate Gyrus anterior division | Salience |
| Right Paracingulate Gyrus | Salience |
| Right Cingulate Gyrus anterior division | Salience |
| Right Frontal Pole | Salience |
| Right Cingulate Gyrus posterior division | Memory retrieval |
| Right Thalamus | Subcortical |
| Left Thalamus | Subcortical |
| Brain-Stem | Subcortical |
| Left Putamen | Subcortical |
| Left Caudate | Subcortical |
| Right Putamen | Subcortical |
| Right Pallidum | Subcortical |
| Left Postcentral Gyrus | Subcortical |
| Left Middle Temporal Gyrus temporooccipital part | Ventral attention |
| Left Planum Temporale | Ventral attention |
| Right Superior Temporal Gyrus posterior division | Ventral attention |
| Right Middle Temporal Gyrus posterior division | Ventral attention |
| Right Middle Temporal Gyrus temporooccipital part | Ventral attention |
| Right Inferior Frontal Gyrus pars triangularis | Ventral attention |
| Left Frontal Operculum Cortex | Ventral attention |
| Left VI | Cerebellar |
| Left VI | Cerebellar |
| Right VI | Cerebellar |
| Vermis VI | Cerebellar |
| Right Temporal Fusiform Cortex posterior division | Uncertain |
| Left Temporal Fusiform Cortex anterior division | Uncertain |
| Right Inferior Temporal Gyrus anterior division | Uncertain |
| Left Inferior Temporal Gyrus anterior division | Uncertain |
| Right Lateral Occipital Cortex superior division | Dorsal attention |
| Left Lateral Occipital Cortex inferior division | Dorsal attention |
| Left Inferior Temporal Gyrus temporooccipital part | Uncertain |
| Right Temporal Occipital Fusiform Cortex | Uncertain |
| Right Lateral Occipital Cortex superior division | Dorsal attention |
| Right Middle Temporal Gyrus temporooccipital part | Dorsal attention |
| Right Lateral Occipital Cortex superior division | Dorsal attention |
| Left Superior Parietal Lobule | Dorsal attention |
| Left Lateral Occipital Cortex superior division | Dorsal attention |
| Left Middle Frontal Gyrus | Dorsal attention |
| Left Inferior Temporal Gyrus temporooccipital part | Dorsal attention |
| Left Lateral Occipital Cortex superior division | Dorsal attention |
| Right Precentral Gyrus | Dorsal attention |

**Table S5:**
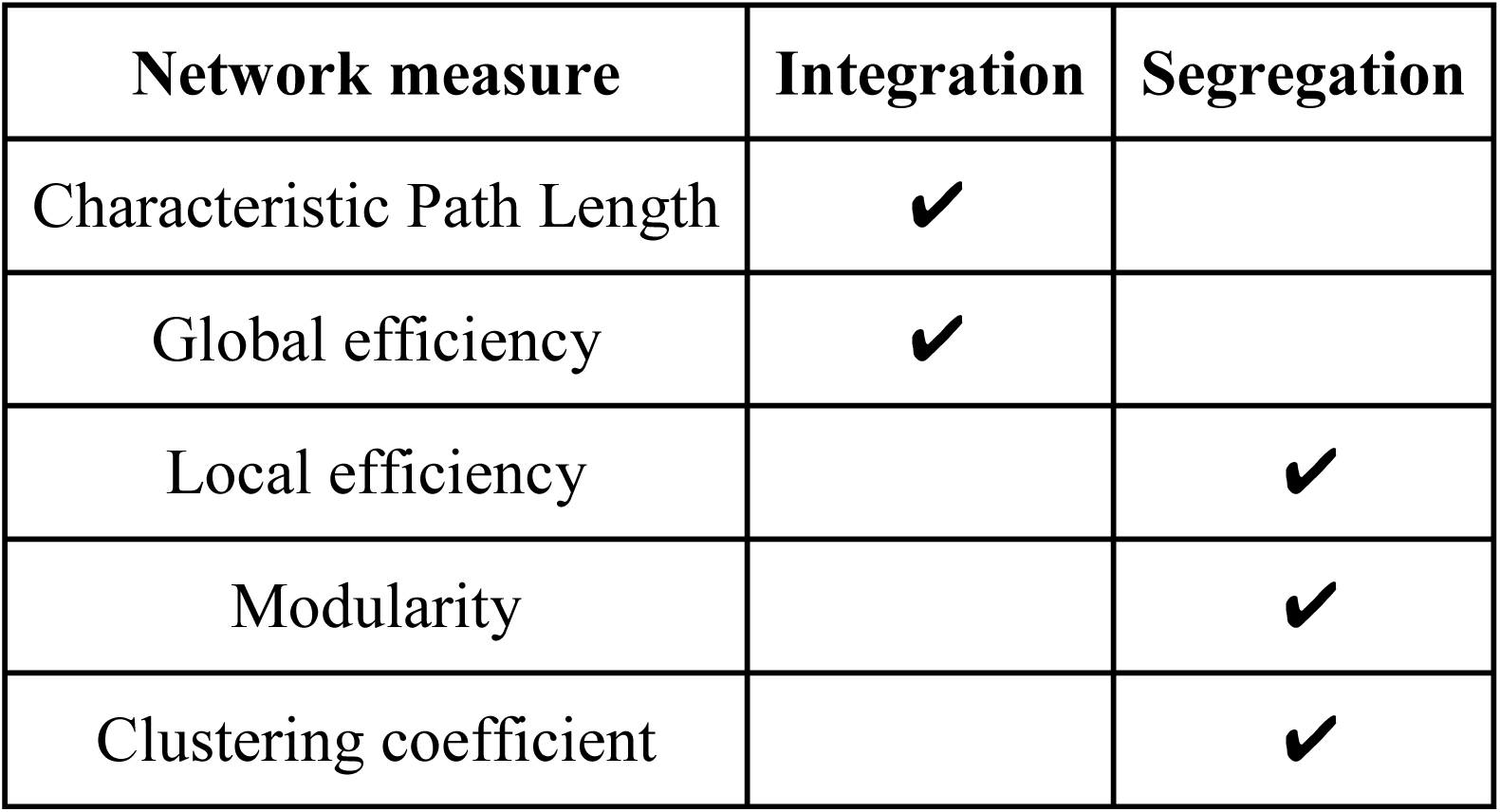
Network measures classified as measures of integration and segregation.

### Supplementary Figures

**Fig. S1:**
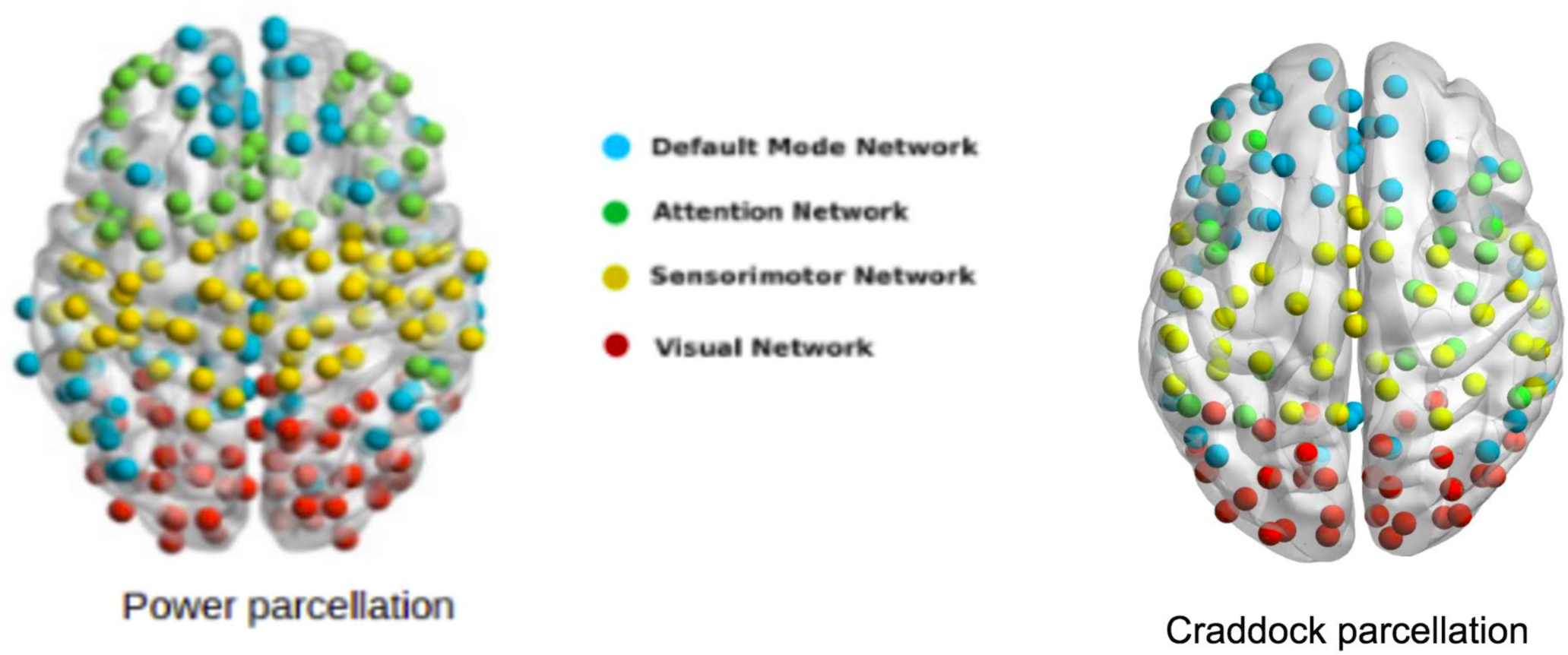
Communities identified in UCLA (Power parcellation) and ABIDE (Craddock parcellation) datasets in TD cohorts. The region of interests belonging to four distinct modules in the brain network: Default Mode, Sensorimotor, Attention and Visual are shown.

**Fig. S2:**
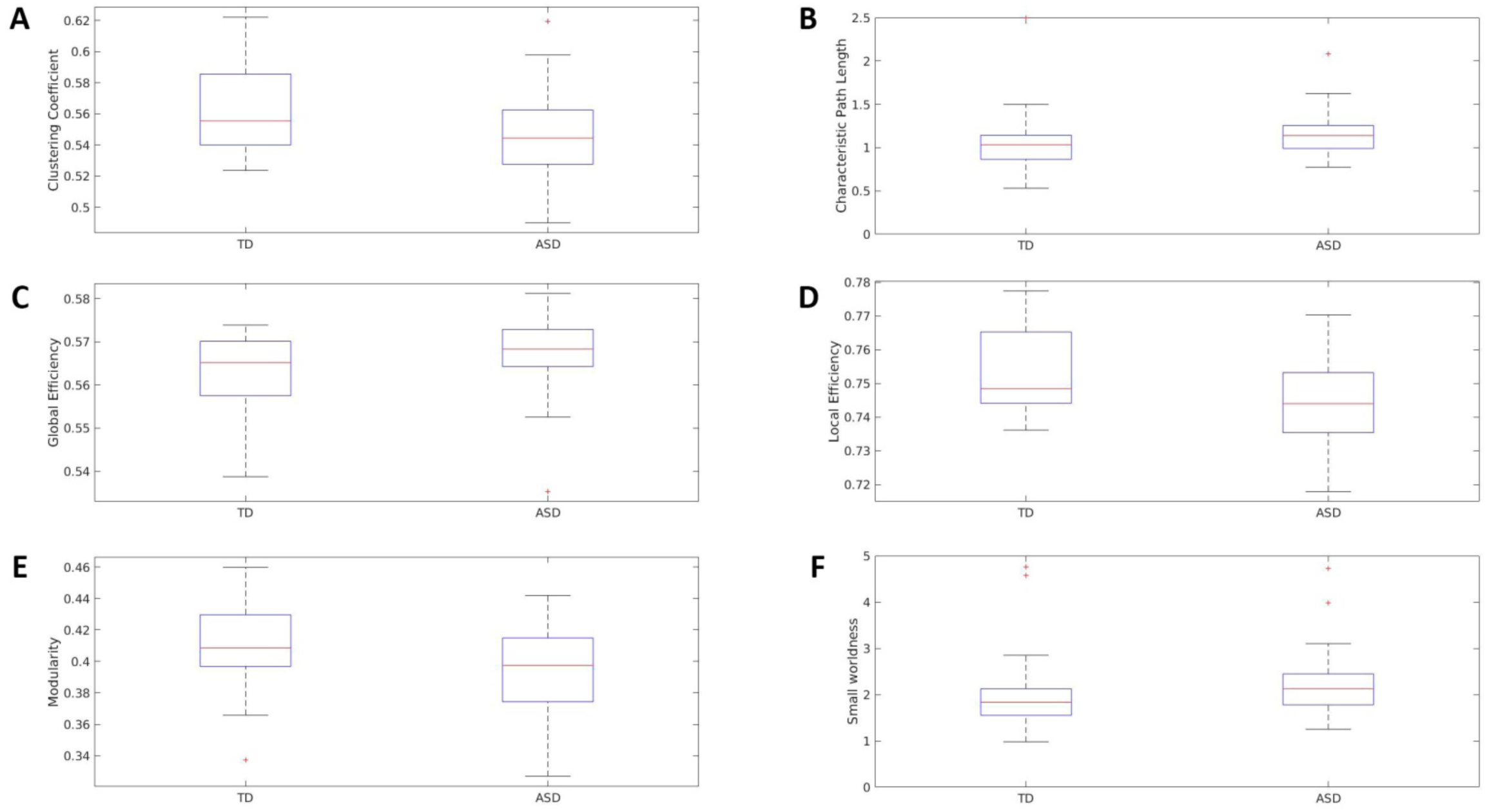
Differences in the network measures of functional integration and segregation between TD and ASD for UCLA database.

**Fig. S3:**
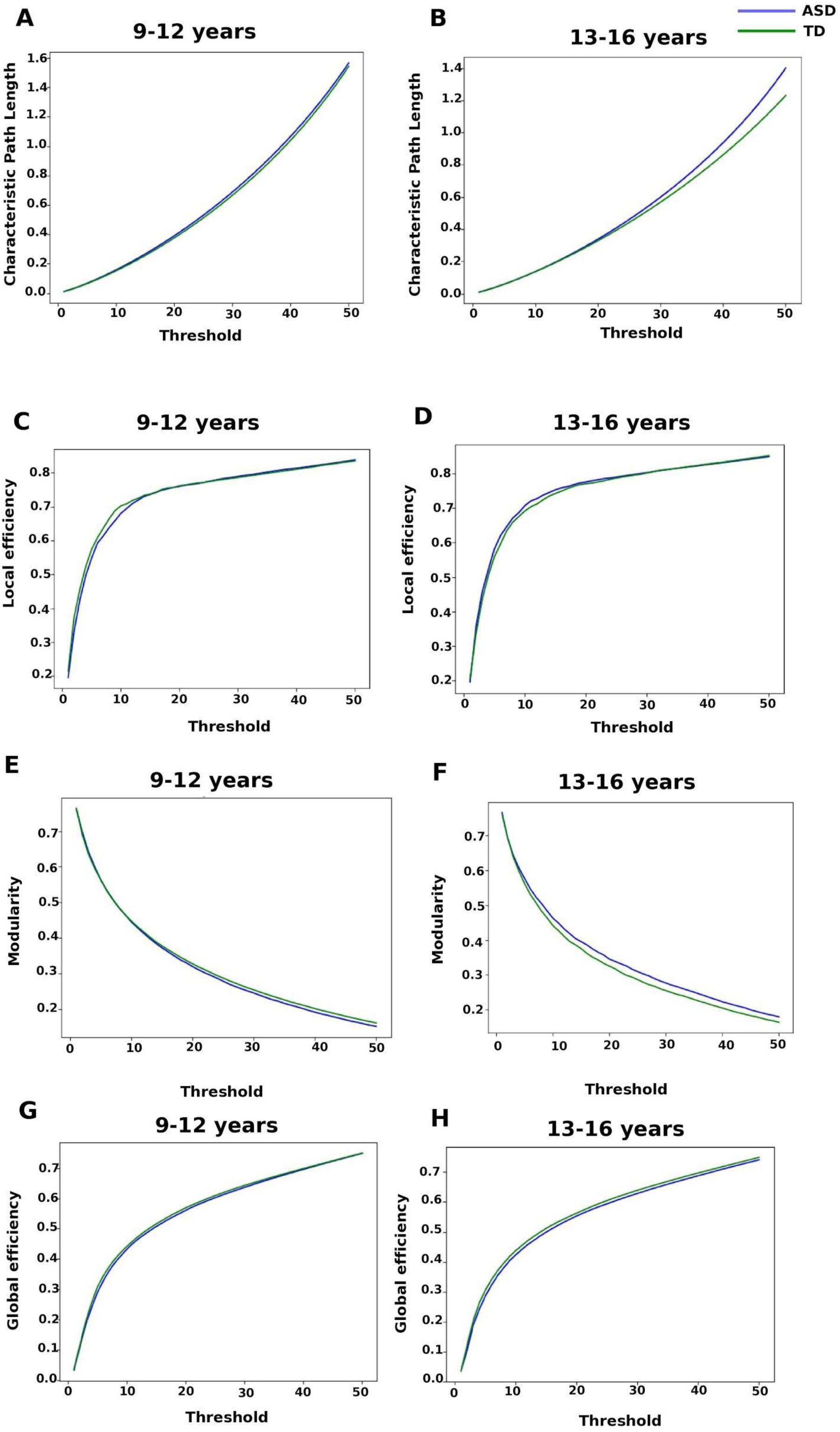
Sensitivity of Network measures to the range of thresholds with respect to age and disease. **(A-B)** Characteristic path length (CPL), **(C-D)** Local efficiency, **(E-F)** Modularity, **(G-H)** Global efficiency.

